# Species limits in butterflies (Lepidoptera: Nymphalidae): Reconciling classical taxonomy with the multispecies coalescent

**DOI:** 10.1101/451039

**Authors:** Pável Matos-Maraví, Niklas Wahlberg, Alexandre Antonelli, Carla M. Penz

## Abstract

Species delimitation is at the core of biological sciences. During the last decade, molecular-based approaches have advanced the field by providing additional sources of evidence to classical, morphology-based taxonomy. However, taxonomy has not yet fully embraced molecular species delimitation beyond threshold-based, single-gene approaches, and taxonomic knowledge is not commonly integrated to multi-locus species delimitation models. Here we aim to bridge empirical data (taxonomic and genetic) with recently developed coalescent-based species delimitation approaches. We use the multispecies coalescent model as implemented in two Bayesian methods (DISSECT/STACEY and BP&P) to infer species hypotheses. In both cases, we account for phylogenetic uncertainty (by not using any guide tree) and taxonomic uncertainty (by measuring the impact of using or not a priori taxonomic assignment to specimens). We focus on an entire Neotropical tribe of butterflies, the Haeterini (Nymphalidae: Satyrinae). We contrast divergent taxonomic opinion—splitting, lumping and misclassifying species—in the light of different phenotypic classifications proposed to date. Our results provide a solid background for the recognition of 22 species. The synergistic approach presented here overcomes limitations in both traditional taxonomy (e.g. by recognizing cryptic species) and molecular-based methods (e.g. by recognizing structured populations, and not raise them to species). Our framework provides a step forward towards standardization and increasing reproducibility of species delimitations.

## INTRODUCTION

Traditionally, taxonomic delimitations have relied on diagnostic phenotypic characters to classify distinct populations into species and subspecies (hereafter, the ‘traditional taxonomic approach’). More recently, coalescent-based methods that quantify reproductive isolation using genetic data have been proposed as a means to calculate the probability of speciation (hereafter, the ‘coalescent approach’; e.g. Knowles & Carstens, 2007; Yang & Rannala, 2010; Fujita *et al.*, 2012). Both approaches do not necessarily agree in their species hypotheses because their scopes are centered on different sections of the speciation continuum; while traditional taxonomy primarily depends on the evolution of informative and consistent phenotypic characters, the coalescent approach is guided by any gene flow reduction that could be associated with the onset of reproductive isolation. As a consequence, phenotype-based delimitation may not identify cryptic and incipient species (i.e. genetically divergent lineages embarked in the process of speciation; Rosindell *et al.*, 2010) whereas coalescent-based delimitation may simply reveal population structure (i.e. subpopulations with a long non-breeding history) (Sukumaran & Knowles, 2017). Despite the paramount importance of delimiting species for multiple disciplines and practices in science and society (e.g. ecology, evolution, conservation biology, among others), it still remains unclear how to reconcile conflicts between traditional taxonomy and the coalescent approach while taking into account their respective benefits and limitations.

Taxonomists have not fully embraced the recent developments in molecular species delimitation beyond threshold-based, single-gene approaches including DNA barcoding based on COI proposed as a fast approach to delimit species (Hebert *et al*., 2004), and delimitation methods based on single markers (Pons *et al*., 2006; Puillandre *et al*., 2012). End-users of the coalescent approach, on the other hand, do not usually incorporate taxonomic knowledge to inform their models (e.g. Leaché & Fujita, 2010; Olave *et al.*, 2014; but see Aydin *et al.*, 2014; Jones *et al.*, 2015). In a Bayesian framework, taxonomic information could be explicitly acknowledged in the form of prior distributions, and thus alternative species hypotheses can be statistically weighed. Indeed, the latest Bayesian implementations, which include the multispecies coalescent model (MSC; Degnan & Rosenberg, 2009), can accommodate parameters that control the number of species in a dataset, their divergence times and ancestral population sizes, all in a single probabilistic framework. These properties among others, such as the recognition of genealogical incongruence and incomplete lineage sorting, arguably make MSC-based methods more biologically realistic than threshold-based molecular species delimitation (Knowles & Carstens, 2007; Fujita *et al.*, 2012). However, it remains unexplored how divergent taxonomic opinions affect species delimitations, when these opinions (such as “splitters” *vs*. “lumpers”) are translated into prior distributions for molecular species delimitation analyses.

Here we aim to reconcile genetic and taxonomic data using species delimitation models that estimate coalescence and species divergence in a fully Bayesian framework. We study the butterflies classified in the tribe Haeterini (Nymphalidae: Satyrinae), insects that exclusively inhabit tropical rainforests in southern Mexico, Central and South America. Transparent wings are the most obvious characteristic of this group, an attribute shared by four out of five genera within the tribe—*Pierella* being the exception by having full scale-cover (Fig. 1).

**Figure 1:**
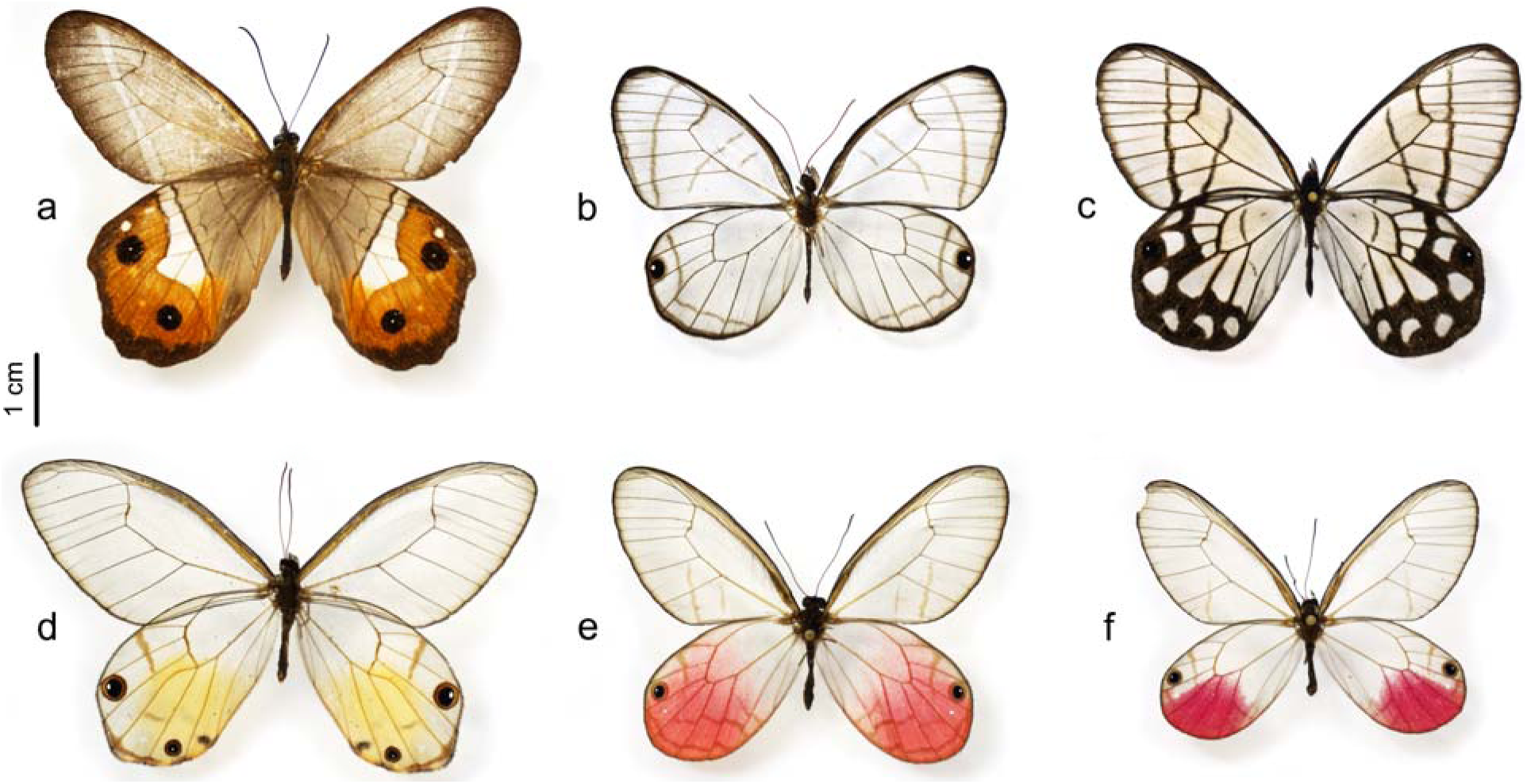
Representatives of the tribe Haeterini. a) *Pierella nereis* (Brazil, Minas Gerais, Santa Barbara; Milwaukee Public Museum), b) *Dulcedo polita* (Costa Rica, Sarapiqui, Tirimbina Biological Station; Phil DeVries Collection), c) *Pseudohaetera mimica* (Peru, Junin, Satipo; Natural History Museum of Los Angeles County), d) *Haetera piera* (Ecuador, Napo, Garza Cocha; Phil DeVries Collection), e) *Cithaerias cliftoni* (Ecuador, Oriente; Natural History Museum of Los Angeles County), f) *Cithaerias aurora tambopata* (Peru, Madre de Dios, Pakitza, Manu National Reserve; Smithsonian Institution).

Taxonomic work on this charismatic group involves numerous researchers over the last 150 years (e.g. Herrich-Schäffer, 1864; Weymer, 1910; Miller, 1968; Constantino, 1995; Lamas, 1997; Penz *et al.*, 2014; Paluch *et al.*, 2015; Willmott, 2015; Zacca *et al.*, 2016). Haeterini is a monophyletic tribe (Wahlberg *et al*., 2009; Chazot *et al*., 2019), and current taxonomic understanding (Lamas, 2004; Penz *et al.*, 2014; Willmott, 2015; Zacca *et al.*, 2016) is that Haeterini consists of 29 described species and 39 subspecies. However, these estimates remain contentious because published taxonomic studies have mostly focused on smaller groups within Haeterini and have relied on different morphological character systems (wing coloration, genitalia shape or male androconial organs), and have not taken into account variation in DNA sequence data. The taxonomic opinions among these authors reflect the well-recognized subjectivity of species-level taxonomic work, even though Haeterini has a comprehensive taxonomic knowledge as compared to most other tropical insects. This offers the opportunity to quantify taxonomic opinion and definitions among authors, as a preliminary step for standardization.

Specifically, we aim to test alternative species delimitation hypotheses and to evaluate the following two interconnected expectations:

1. Lineage delimitation using the multispecies coalescent model (MSC) will recover the taxonomic subspecies in Haeterini. Butterfly subspecies are commonly described based on parapatric or allopatric geographical distributions, and reduced gene flow could facilitate phenotypic divergence. If phenotype divergence resulted from genetic differentiation, then any reproductive isolation among subspecies will be recovered by the MSC.

2. Taxonomic knowledge modeled as prior distributions will group specimens into MSC Clusters (MSCC). However, regardless of prior distribution, the MSC will standardize species delimitation because divergent taxonomic opinion among authors, if any, will be evident in the phylogeny of Haeterini. Therefore, the MSC will inform on whether the use of different morphological characters (wing coloration, genitalia shape or male androconial organs) might favor narrower or broader species definitions in Haeterini butterflies.

## MATERIALS AND METHODS

### Taxon sampling and molecular dataset

Haeterini butterflies were collected by the authors and collaborators throughout most of the geographical range of the tribe, including localities from northern Costa Rica to southeastern Brazil. Specimens were identified to the species and subspecies level following published taxonomic revisions (Constantino, 1995; Lamas, 1997; Penz *et al.*, 2014; Paluch *et al.*, 2015; Willmott, 2015; Zacca *et al.*, 2016) and comparing wing morphology to type specimen photographs at https://www.butterfliesofamerica.com (last accessed August 2018). We sampled all five Haeterini genera, including 18 of 29 currently valid species, and 18 of 39 subspecies (Lamas, 2004; Penz *et al.*, 2014; Paluch *et al.*, 2015; Willmott, 2015; Zacca *et al.*, 2016). Note that several subspecies were not represented in our sample because they are only known from type collections (hampering their access for genetic studies) or have rarely been collected in recent years.

We followed standard lab protocols (Wahlberg & Wheat, 2008) to sequence 6 gene fragments from 63 specimens: the mitochondrial locus COI (1,475 bp) and the nuclear loci CAD (850 bp), EF1α (1,240 bp), GAPDH (691 bp), RpS5 (617 bp), and *wingless* (400 bp). Sanger sequencing was conducted by the company Macrogen (South Korea), and sequence quality control and DNA alignments were carried out using the program Geneious R7. We retrieved from GenBank the DNA sequences of sixteen species classified in the subfamily Satyrinae in order to root the phylogeny of Haeterini. All DNA sequences were deposited in GenBank (accession numbers MH802134–MH802346, see Table 1) and the DNA alignments used in this study were archived in TreeBASE (ID 23439). Detailed specimen voucher information can be found in Table 1.

**Table 1:** Voucher locality information and associated genetic data. All Haeterini specimens were identified to the species and subspecies level based on the most recent taxonomic revisions (Constantino 1995; Lamas 1997; Penz et al. 2014; Paluch et al. 2015; Willmott 2015; Zacca et al. 2016). GenBank accession numbers for each of the sequenced locus are presented for every specimen.

| Code | Genus | Species | Subspecies | Country | Locality | CAD | COI | EF1a | GAPDH | RpS5 | wingless |
| --- | --- | --- | --- | --- | --- | --- | --- | --- | --- | --- | --- |
| EW1-1 | Pararge | aegeria |  | FRANCE | Carcassonne, 2 km S Trassonel | EU141293 | DQ176379 | DQ338913 | EU141476 | EU141372 | DQ338620 |
| NW121-17 | Lethe | minerva |  | INDONESIA | Bali | EU141309 | DQ338768 | DQ338909 | EU141492 | EU141387 | DQ338616 |
| EW10-5 | Bicyclus | anyana |  | ZIMBABWE | Harare | EU141295 | AY218238 | AY218258 | EU141478 | EU141374 | AY218276 |
| CP06-89 | Oressinoma | sorata |  | PERU | Pasco, Oxapampa | MH802140 | GQ357209 | GQ357278 | GQ357440 | GQ357570 | GQ357342 |
| EW25-17 | Orsotriaena | medus |  | BANGLADESH | Sylhet Div., Lowacherra Forest | - | DQ338766 | DQ338906 | EU528405 | EU528453 | DQ338633 |
| NW108-3 | Taygetis | virgilia |  | BRAZIL | SP, Campinas, Santa Genebra | EU141305 | DQ338812 | DQ338958 | EU141487 | EU141383 | DQ338683 |
| PM07-05 | Calisto | smintheus |  | CUBA | Alrededores de La Platica | JN881807 | JN881904 | JN881778 | JN881827 | JN881845 | JN881870 |
| CP03-63 | Manerebia | cyclopina |  | PERU | Junín, Quebrada Siete Jeringas | - | DQ338785 | DQ338928 | EU528397 | EU528443 | GQ864477 |
| NW162-21 | Satyrus | actaea |  | FRANCE | Aude, Villegly | GQ864709 | GQ864807 | GQ864901 | GQ865030 | GQ865494 | GQ864495 |
| EW24-7 | Erebia | oeme |  | FRANCE | Languedoc, Ariege 09, Ustou | EU141296 | DQ338780 | DQ338923 | EU141479 | EU141375 | DQ338640 |
| CP13-01 | Aeropetes | tulbaghia |  | S. AFRICA | Mpumalanga Verloren Valei | - | DQ338579 | DQ338907 | EU528381 | EU528419 | DQ338634 |
| NW66-6 | Melanitis | leda |  | AUSTRALIA | Queensland, Cairns | EU141330 | AY090207 | AY090173 | EU141508 | EU141408 | AY090140 |
| NW122-21 | Brassolis | sophorae |  | BRAZIL | SP, Campinas | - | EU528314 | EU528291 | GQ357384 | EU528425 | EU528270 |
| NW66-5 | Morpho | helenor |  | - | London Pupae Supplies | EU141329 | AY090210 | AY090176 | EU141507 | EU141407 | AY090143 |
| NW121-20 | Elymnias | casiphone |  | INDONESIA | Bali | EU141310 | DQ338760 | DQ338900 | - | EU141388 | DQ338627 |
| NW101-2 | Zeuxidia | dorhni |  | INDONESIA | Java, Bandung | - | DQ338752 | DQ338892 | EU528417 | EU528471 | DQ338609 |
| PM14-21 | Cithaerias | andromeda | bandusia | BRAZIL | MT, Alta Floresta | KJ145993 | KJ145988 | KJ145996 | KJ146001 | KJ146004 | KJ146008 |
| NW92-1 | Cithaerias | aurora | aurora | PERU | Loreto, Río Paiwa | - | MH802187 | - | MH802251 | MH802286 | MH802332 |
| NW93-1 | Cithaerias | aurora | aurora | PERU | Loreto, Río Paiwa | - | DQ338756 | DQ338896 | MH802252 | MH802287 | DQ338613 |
| CP01-52 | Cithaerias | aurora | tambopata | PERU | Tambopata Research Center | MH802134 | MH802157 | MH802205 | MH802239 | MH802258 | MH802304 |
| LEP-08919 | Cithaerias | cliftoni |  | ECUADOR | Pastaza, Kapawi lodge | MH802148 | MH802174 | MH802218 | MH802246 | MH802276 | MH802320 |
| LEP-10406 | Cithaerias | cliftoni |  | ECUADOR | Sucumbios, Cuyabeno lodge | KM013123 | KM012939 | KM012990 | KM013275 | KM013174 | KM013055 |
| PM14-17 | Cithaerias | cliftoni |  | ECUADOR | Sucumbios, Garza Cocha | - | MH802196 | - | - | MH802295 | MH802341 |
| PM24-06 | Cithaerias | pireta | pireta | COSTA RICA | Sarapiquí, Agrícola Sofia, nr. Tirimbina | - | - | MH802238 | - | MH802299 | MH802345 |
| LEP-09925 | Cithaerias | pireta | pireta | ECUADOR | Carchi, Finca San Francisco | MH802154 | MH802182 | MH802225 | MH802249 | MH802282 | MH802327 |
| LEP-09926 | Cithaerias | pireta | pireta | ECUADOR | Pichincha, km. 20 Pacto-Guayabillas Rd. | KM013135 | KM012940 | KM013007 | KM013282 | KM013181 | KM013069 |
| BCI86376 | Cithaerias | pireta | pireta | PANAMA | BCI | - | HM406591 | MH802203 | - | MH802256 | MH802302 |
| LEP-14256 | Cithaerias | pyropina | julia | ECUADOR | Morona Santiago | - | MH802183 | MH802226 | - | MH802283 | MH802328 |
| LEP-14257 | Cithaerias | pyropina | julia | ECUADOR | Morona Santiago | - | MH802184 | MH802227 | - | MH802284 | MH802329 |
| PM0095 | Cithaerias | pyropina | pyropina | PERU | Pasco, Cañón de Huancabamba | - | MH802192 | MH802232 | - | MH802291 | MH802337 |
| PM11-05 | Dulcedo | polita |  | COSTA RICA | Heredia, Tirimbina | - | KJ145990 | KJ145998 | KJ146002 | KJ146006 | KJ146010 |
| LEP-14258 | Dulcedo | polita |  | ECUADOR | Esmeraldas, Río Chuchubi | - | MH802185 | MH802228 | MH802250 | - | MH802330 |
| FLMNH284117 | Haetera | macleaniania |  | ECUADOR | Esmeraldas, Finca Cypis | - | MH802165 | MH802209 | MH802242 | MH802267 | MH802311 |
| LEP-04409 | Haetera | piera | negra | ECUADOR | Morona Santiago, nr. Yaupi | MH802142 | MH802167 | MH802211 | MH802244 | MH802269 | MH802313 |
| LEP-08916 | Haetera | piera | negra | ECUADOR | Pastaza, Kapawi lodge | MH802147 | MH802173 | MH802217 | MH802245 | MH802275 | MH802319 |
| PM0091 | Haetera | piera | negra | PERU | Pasco, Cañón de Huancabamba | - | MH802191 | MH802231 | MH802253 | MH802290 | MH802336 |
| CP01-84 | Haetera | piera | pakitza | PERU | Tambopata Research Center | EU141292 | DQ018959 | DQ018926 | EU141475 | EU141371 | DQ018897 |
| PM14-20 | Pierella | chalybaea |  | BRAZIL | MT, Alta Floresta | KM013148 | KM012978 | KM013025 | - | KM013197 | KM013087 |
| LEP-04597 | Pierella | chalybaea |  | ECUADOR | Orellana, Napo Wildlife Center | - | MH802168 | MH802212 | - | MH802270 | MH802314 |
| LEP-08934 | Pierella | chalybaea |  | ECUADOR | Pastaza, Kapawi lodge | MH802151 | MH802178 | MH802221 | - | MH802279 | MH802323 |
| PM14-11 | Pierella | chalybaea |  | ECUADOR | Sucumbios, Garza Cocha | MH802155 | MH802193 | MH802233 | - | MH802292 | MH802338 |
| PM24-01 | Pierella | helvina | incanescens | COSTA RICA | Sarapiquí, Finca Starke | - | MH802197 | - | - | - | - |
| PM24-02 | Pierella | helvina | incanescens | COSTA RICA | Heredia, Tirimbina | - | MH802198 | MH802235 | - | MH802296 | MH802342 |
| PM24-03 | Pierella | helvina | incanescens | COSTA RICA | Heredia, Tirimbina | - | MH802199 | MH802236 | MH802254 | MH802297 | MH802343 |
| FLMNH146191 | Pierella | helvina | ocreata | ECUADOR | Esmeraldas, Río Chuchubi | - | MH802164 | MH802208 | - | - | MH802310 |
| LEP55828 | Pierella | helvina | ocreata | ECUADOR | Esmeraldas, San Francisco Ridge | - | MH802166 | MH802210 | MH802243 | MH802268 | MH802312 |
| BCI80049 | Pierella | helvina | ocreata | PANAMA | BCI | - | MH802156 | MH802202 | - | MH802255 | MH802301 |
| CP01-69 | Pierella | hortona | albofasciata | PERU | Tambopata Research Center | MH802135 | - | MH802206 | - | MH802259 | - |
| LEP-08928 | Pierella | hortona | hortona | ECUADOR | Pastaza, Kapawi lodge | MH802150 | MH802177 | MH802220 | - | MH802278 | MH802322 |
| CP05-34 | Pierella | hortona | hortona | PERU | Paraíso, Río Momón | MH802139 | - | - | - | MH802265 | MH802308 |
| CP02-85 | Pierella | hyceta | ceryce | PERU | Junín, Aldea | MH802136 | MH802158 | - | - | MH802260 | MH802305 |
| CP04-47 | Pierella | hyceta | ceryce | PERU | Junín, Río Colorado, Quebrada Perla | MH802138 | MH802160 | - | - | MH802263 | MH802307 |
| LEP-08925 | Pierella | hyceta | hyceta | ECUADOR | Zamora-Chinchipe, San Roque, Ridge E | KJ145994 | KJ145989 | KJ145997 | - | KJ146005 | KJ146009 |
| LEP-08926 | Pierella | hyceta | hyceta | ECUADOR |  | - | MH802176 | - | - | - | - |
| CP03-60 | Pierella | hyceta | hyceta | PERU | Yanachaga-Chemillén | MH802137 | MH802159 | - | - | MH802261 | MH802306 |
| PM0089 | Pierella | hyceta | hyceta | PERU | Pasco, Cañón de Huancabamba | - | MH802189 | MH802230 | - | MH802289 | MH802334 |
| PM0090 | Pierella | hyceta | hyceta | PERU | Pasco, Cañón de Huancabamba | - | MH802190 | - | - | - | MH802335 |
| SnPe05 | Pierella | hyceta | hyceta | PERU | Cuzco, San Pedro | - | MH802201 | - | - | MH802300 | MH802346 |
| NW108-2 | Pierella | keithbrowni |  | BRAZIL | SP, Ilha do Cardoso, Cananéia | - | MH802186 | MH802229 | - | MH802285 | MH802331 |
| LEP-08908 | Pierella | lena | brasiliensis (browni) | ECUADOR | Pastaza, Kapawi lodge | MH802144 | MH802170 | MH802214 | - | MH802272 | MH802316 |
| LEP-08911 | Pierella | lena | brasiliensis | ECUADOR | Morona-Santiago, Wachirpas airfield | MH802145 | MH802171 | MH802215 | - | MH802273 | MH802317 |
| LEP-08912 | Pierella | lena | brasiliensis | ECUADOR | Pastaza, Kapawi lodge | MH802146 | MH802172 | MH802216 | - | MH802274 | MH802318 |
| NW93-2 | Pierella | lena | brasiliensis | PERU | Loreto, Río Paiwa | - | DQ338757 | DQ338897 | - | MH802288 | DQ338614 |
| PM14-19 | Pierella | lena | lena | BRAZIL | MT, Alta Floresta | KM013139 | KM012979 | KM013011 | - | KM013185 | KM013073 |
| PM14-13 | Pierella | lena | lena | ECUADOR | Sucumbios, Garza Cocha | - | MH802195 | - | - | MH802294 | MH802340 |
| LEP-04613 | Pierella | lucia |  | ECUADOR | Orellana, Napo Wildlife Center | MH802143 | MH802169 | MH802213 | - | MH802271 | MH802315 |
| LEP-08935 | Pierella | lucia |  | ECUADOR | Pastaza, Kapawi lodge | MH802152 | MH802179 | MH802222 | - | MH802280 | MH802324 |
| PM14-12 | Pierella | lucia |  | ECUADOR | Sucumbios, Garza Cocha | - | MH802194 | MH802234 | - | MH802293 | MH802339 |
| CP05-11 | Pierella | lucia |  | PERU | Cordillera del Cóndor | - | MH802162 | - | - | - | - |
| LEP-09797 | Pierella | luna | lesbia | ECUADOR | Manabi, above Camarones, Pedernales-Jama Rd. | MH802153 | MH802181 | MH802224 | - | MH802281 | MH802326 |
| LEP-09921 | Pierella | luna | lesbia | ECUADOR | Esmeraldas, Tundaloma lodge | KM013149 | KM012980 | KM013026 | - | KM013198 | KM013088 |
| PM24-04 | Pierella | luna | luna | COSTA RICA | Heredia, Tirimbina | - | MH802200 | MH802237 | - | MH802298 | MH802344 |
| BCI90713 | Pierella | luna | luna | PANAMA | Argos Plantations | - | HM406585 | MH802204 | - | MH802257 | MH802303 |
| CP14-08 | Pierella | nereis |  | BRAZIL | SP, São Luiz do Paraitingo | MH802141 | MH802163 | MH802207 | - | MH802266 | MH802309 |
| LEP-08924 | Pseudohaetera | hypaesia |  | ECUADOR | Zamora-Chinchipe, Quebrada Guayzimi (close to La Merced) | MH802149 | MH802175 | MH802219 | MH802247 | MH802277 | MH802321 |
| LEP-09562 | Pseudohaetera | hypaesia |  | ECUADOR | Morona Santiago, El Boliche | - | MH802180 | MH802223 | MH802248 | - | MH802325 |
| CP03-99 | Pseudohaetera | hypaesia |  | PERU | Junín, Quebrada Siete Jeringas | - | DQ338758 | DQ338898 | MH802240 | MH802262 | DQ338625 |
| CP04-57 | Pseudohaetera | hypaesia |  | PERU | Junín, Mina Pichita | - | MH802161 | - | MH802241 | MH802264 | - |
| PM0062 | Pseudohaetera | hypaesia |  | PERU | Junín, San Ramón, Nueva Italia | - | MH802188 | - | - | - | MH802333 |

### Phylogenetic analyses

We inferred phylogenies using single-gene datasets partitioned by codon position to rule out any tree topology conflict due to contamination. In addition, in order to test the monophyly of each genus, we inferred a phylogeny using the concatenated multi-locus dataset consisting of 6 genetic markers that proved to be phylogenetically informative (Table 2), and 63 Haeterini specimens and 16 outgroup taxa. We used PartitionFinder v2.1.1 (Lanfear *et al.*, 2017) to estimate the best-fit partitioning strategy for the concatenated dataset using 18 data blocks (each codon position separately for each gene region) and the following settings: branchlengths = linked (higher likelihood than the unlinked option), models = mrbayes, model_selection = bic and search = greedy. All phylogenetic analyses using single-gene and concatenated datasets were carried out using MrBayes v3.2.3 (Ronquist *et al.*, 2012) through the CIPRES portal (Miller *et al.*, 2010). We used the reversible-jump Markov chain Monte Carlo approach (rjMCMC) to allow moving across nucleotide substitution schemes (nst = mixed) with different rate variation across sites (+I and +Γ). This approach integrated the uncertainty of substitution models that fit the data with the inference of tree topology and branch lengths. Two independent analyses and four chains, one cold and three heated, were run for 10 million cycles and sampled every 1,000 cycles, discarding the first 25% sampled parameters as burn-in. We evaluated convergence using the average standard deviation of split frequencies (< 0.005), potential scale reduction factor (~ 1.000), estimated sample sizes (ESS > 200), and by inspection of stationary distribution of log-probabilities in both independent runs. We summarized the 7,500 sampled trees using the 50% majority-rule consensus method.

**Table 2:** Characteristics of the molecular dataset used in this study, including gene and specimen coverage, GC content and the number of variable sites.

| Genes | Specimens | Length (bp) | Variable sites | Missing data (%) | GC content (%) |
| --- | --- | --- | --- | --- | --- |
| CAD | 29 (46%) | 850 | 157 | 25.8 | 33.5 |
| COI | 60 (95%) | 1475 | 474 | 26.5 | 29.5 |
| EF1α | 49 (78%) | 1240 | 230 | 17.5 | 48.6 |
| GAPDH | 21 (33%) | 691 | 129 | 2.6 | 45.4 |
| RpS5 | 55 (87%) | 617 | 124 | 1.9 | 45.2 |
| *wingless* | 58 (92%) | 412 | 96 | 9.3 | 58.6 |
| TOTAL | 63 | 5285 | 1210 | 39.2 | 41.4 |

### Molecular species delimitation

Once we had confidence on the monophyly of each genus and a notion of the phylogenetic relationships among the 63 specimens, we carried out molecular species delimitation under the MSC framework. Although the inference of phylogenetic relationships is not a pre-requisite for subsequent delimitation analyses, it becomes informative when large phylogenies need to be divided into smaller well-supported subclades, and analyzed separately due to computational limitation (see BP&P analyses below). We used a comprehensive taxon sampling of Haeterini, a multi-locus dataset, and two popular Bayesian implementations of the MSC, namely STACEY (Jones, 2017) and BP&P (Yang, 2015):

#### STACEY

We used DISSECT (Jones *et al.*, 2015) which is a taxonomic assignment-free Bayesian method for grouping individuals into multispecies coalescent clusters (MSCC). The method is implemented in the STACEY v1.2.4 package (Jones, 2017) available in BEAST v2.4.7 (Bouckaert *et al.*, 2014). All gene markers sequenced for this study are likely unlinked in the genome of the butterflies and thus gene trees, substitution and clock models were all treated as unlinked in the analysis. We assigned to the mitochondrial COI locus a gene ploidy of 0.5 and to the remaining nuclear loci a gene ploidy of 2.0 (diploid). Uncorrelated relaxed-clock models were chosen for all loci, and we estimated nuclear clock rates relative to the COI mean clock rate fixed to 1.0. The relative clock mean priors were all log normal (M = 0, S = 1). We used the birth-death-collapse model following Jones *et al.* (2015) with GrowthRate prior as log normal (M = 5, S = 2) and relative DeathRate as uniform in [0, 1], while the popMean prior was set to log normal (M = −7, S = 2).

In STACEY, the discovery of MSCCs relies on two parameters that control node collapsing in the phylogeny, the collapseHeight (ε) and collapseWeight (ω). The parameter ε distinguishes very shallow species divergences (node heights) and should be assigned a small value (Jones *et al.*, 2015), thus, we set ε to 1*e*−4. The parameter ω controls the number of MSCCs and can be used as a proxy for prior taxonomic knowledge. The 1/X distribution of the prior for the number of MSCCs has a mean of 1 + (*n* − 1) × (1 − ω), where *n* is the number of individuals in the dataset. We set ω to 0.73 or 0.59, corresponding to 18 described species and 26 taxa (the 18 sampled subspecies elevated to species), respectively. In addition, we carried out a third analysis that did not take into account prior taxonomic information on the number of species by using a Beta distribution (α = 2, β= 2) as a prior for the parameter ω. The analyses were run four independent times for 200 million cycles each, with parameters sampled every 20,000 cycles. Sampled trees were combined after discarding the first 25% samples as burn-in and checking that ESS values were > 200. MSCCs, their posterior probabilities and pairwise similarity probabilities were obtained using SpeciesDelimitationAnalyser v1.8.0 (Jones *et al.*, 2015) acknowledging ε = 1*e*−4. We chose the clustering with the highest posterior probability (counts) as the working species hypothesis.

#### BP&P

We used the multispecies coalescent model as implemented in BP&P v3.4 (Yang, 2015) to jointly infer species trees and delimit MSCCs, without using any guide tree (Yang & Rannala, 2014; Rannala & Yang, 2017) nor taxonomic assignment to specimens (Yang & Rannala, 2010; Rannala & Yang, 2013). The heredity scalar was set to 1 for all nuclear loci and to 0.25 for the mitochondrial locus. The prior for ancestral population sizes controlled by the parameter θ was assigned the inverse gamma distribution (IG[α, β]), and we evaluated three different scenarios: i) large ancestral population size (IG[3, 0.2]), ii) medium ancestral population size IG[3, 0.1], and iii) small ancestral population size IG[3, 0.02]. These three different scenarios are expected to impact the inferred number of MSCCs, so that a larger ancestral population prior would favor fewer species in the model. Leaché and Fujita (2010) showed that scenario (i) is a conservative combination of parameters, because it favors fewer species; as a counterpart, we have used scenario (iii) with prior mean an order of magnitude lower than in scenario (i). We separately analyzed two Haeterini subclades, namely i) *Pierella* and ii) the remaining genera, to overcome computational limitation in BP&P. We used the inverse gamma distribution for the divergence time of the root in the species trees, which is controlled by the parameter τ_0_. The prior distribution of τ_0_ was diffuse (α = 3) and we specified β = 0.042 for the *Pierella* species tree and β = 0.098 for the remaining Haeterini genera. These values enforced a sequence divergence mean of 2.1% for *Pierella* and 4.9% for the remaining genera, which translate into absolute times of ~7 Ma for *Pierella*’s crown age and ~17 Ma for the remaining genera, assuming a butterfly mutation rate of 2.9 × 10^−9^ (Keightley *et al.*, 2015). The analyses were run two independent times using the rjMCMC algorithm 1 with gamma variable fine-tuning shape α = 2 and mean *m* = 1, each for 500,000 cycles with sampling frequency of 50 and burn-in of the first 10,000 cycles. We chose the species delimitation model with the highest posterior probability as the working species hypothesis.

### Species hypothesis testing and divergence time estimation

We compared the statistical support (model adequacy) for eight species hypotheses in a fully Bayesian framework:

(i) Taxonomic species (18 lineages);

(ii) Taxonomic subspecies elevated to species (26 lineages);

(iii) STACEY’s clusters under ω = 0.73 (accounting for number of taxonomic species; 22 lineages);

(iv) STACEY’s clusters under ω = 0.59 (accounting for number of subspecies elevated to species; 24 lineages);

(v) STACEY’s clusters under ω = Beta (α = 2, β = 2) (non-informative prior on the number of MSCCs; 63 lineages);

(vi) BP&P’s clusters under θ = IG[3, 0.2] (large ancestral population size prior; 21 lineages);

(vii) BP&P’s clusters under θ = IG[3, 0.1] (medium ancestral population size prior; 28 lineages);

(viii) BP&P’s clusters under θ = IG[3, 0.02] (small ancestral population size prior; 55 lineages).

In order to account for incomplete lineage sorting and to avoid any of the pitfalls of using concatenated datasets (Edwards *et al.*, 2016; Bravo *et al.*, 2018), we inferred species tree topology and divergence times using the Bayesian multispecies coalescent method implemented in StarBEAST2 (Ogilvie *et al.*, 2017). We did not use the concatenated-based partition estimated by PartitionFinder in the framework of the multispecies coalescent model because 1) gene trees should be unlinked, 2) each linked codon position within a single locus should share the same clock, and 3) mitochondrial and nuclear loci do not share the same ploidy level. Thus, we used jModelTest v2.1.7 (Darriba *et al.*, 2012) to evaluate the substitution models available in starBEAST2, including or not rate variation among sites (+I and +Γ), for each gene locus. Nucleotide substitution models for each gene locus were chosen on the basis of the Bayesian Information Criterion (BIC). We preliminarily evaluated the fit of two tree models, namely Yule and birth-death, for all 8 species delimitation scenarios using 50 path-sampling steps under thermodynamic integration (Lartillot & Philippe, 2006), each running for 60 million cycles to ensure final ESS values > 200. The Yule tree model had the highest marginal likelihood estimate in all cases and it was preferred over the birth-death model. We therefore report here the path-sampling analyses based on the Yule tree model for the eight species delimitation hypotheses. Other parameters, including gene ploidy, clock models and popMean prior, were set as in the STACEY analyses. The support for each of the eight species delimitation hypotheses was assessed via Bayes factors (*ln*Bf) (Kass & Raftery, 1995) calculated from the posterior tree samples, and we considered *ln*Bf = 2–10 to represent positive but not conclusive support and *ln*Bf > 10 as decisive support for the species hypothesis with the highest marginal likelihood estimated through path sampling.

We must rely on secondary calibrations to date the phylogeny of Haeterini because there are no described fossils assigned to the tribe. Based on a densely sampled, fossil-calibrated butterfly phylogeny (Chazot *et al*., 2019), we constrained the ages of six well-supported nodes that do not belong to Haeterini. We followed a conservative approach by using uniform priors encompassing the 95% highest posterior density (HPD) intervals from (Chazot *et al*., 2019). The constrained nodes included the divergences of:

(i) Brassolini and Morphini to 32–58 Ma;

(ii) Melaniti and Dirini to 23–47 Ma;

(iii) Lethina, Parargina and Mycalesina to 25–44 Ma;

(iv) The crown age of Satyrini to 32–53 Ma;

(v) The crown age of the Satyrini’s subclade encompassing the tribes Pronophilina, Euptychiina, Satyrina, Erebiina, and other closely related subtribes, to 25–43 Ma;

(vi) The crown age of Satyrinae to 41–67 Ma.

Time-calibrated species trees were inferred using BEAST v2.4.7 and the Yule tree prior. Eight independent analyses were carried out to estimate divergence times for the eight species hypotheses described above. Each analysis was run four independent times for 200 million cycles each, with parameters sampled every 20,000 cycles. Trees were summarized in TreeAnnotator (part of BEAST v2.4.7) using the maximum clade credibility method after discarding the first 25% samples as burn-in and merging the four independent runs in LogCombiner (part of BEAST v2.4.7). Convergence among runs was evaluated on the basis of ESS values > 200. DNA alignments and the eight time-calibrated species trees were deposited at Figshare (https://doi.org/10.6084/m9.figshare.7637021.v1). For qualitative evaluation we present “cloudograms” (Figs 2, 3), which are phylogenetic diagrams that reflect topological uncertainty of species trees. Cloudograms were recovered using DensiTree v2.2.5 (Bouckaert, 2010) based on 500 trees from the posterior distribution.

**Figure 2:**
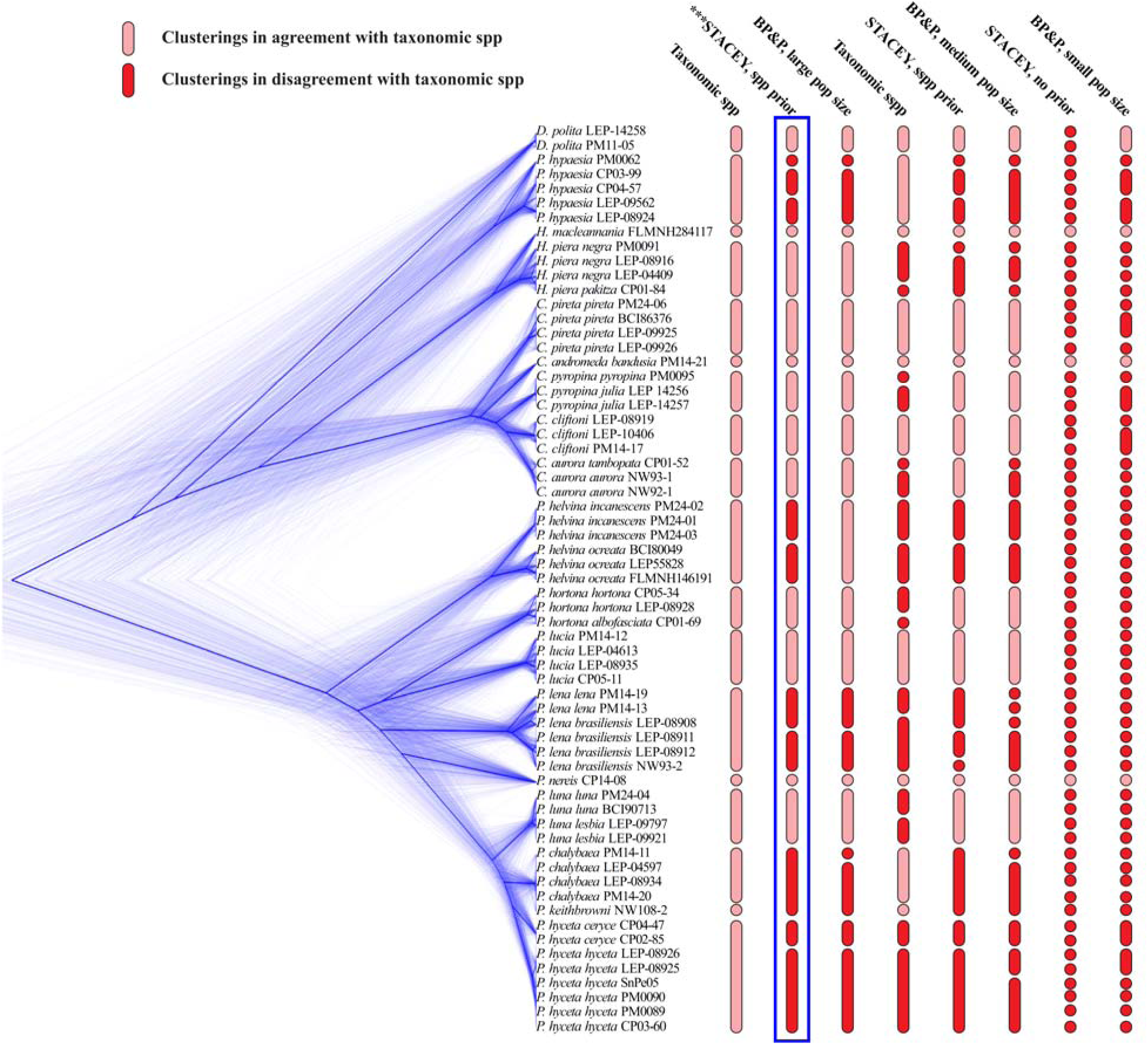
Evaluated species delimitation hypotheses using Bayes factors. The eight scenarios were: Taxonomic species (*spp*, 18 lineages) or subspecies raised to species (*sspp*, 26 lineages), STACEY’s delimited species under prior accounting for number of taxonomic species (*spp prior*, 22 lineages) or number of subspecies raised to species (*sspp prior*, 24 lineages), as well as with prior not informed by taxonomy (*no prior*, 63 lineages), and BP&P’s delimited species under prior for ancestral population size as large (21 lineages), medium (28 lineages), or small (53 lineages). The “cloudogram”, which is a diagram representing phylogenetic uncertainty of the 63 Haeterini specimens, was generated based on 500 posterior trees from STACEY analysis (thicker blue line represents the consensus phylogeny). The delimitation model STACEY under prior accounting for taxonomic species (outlined by a surrounding box) received significant support based on Bayes factors over all other models, and thus is the classification that we propose here.

**Figure 3:**
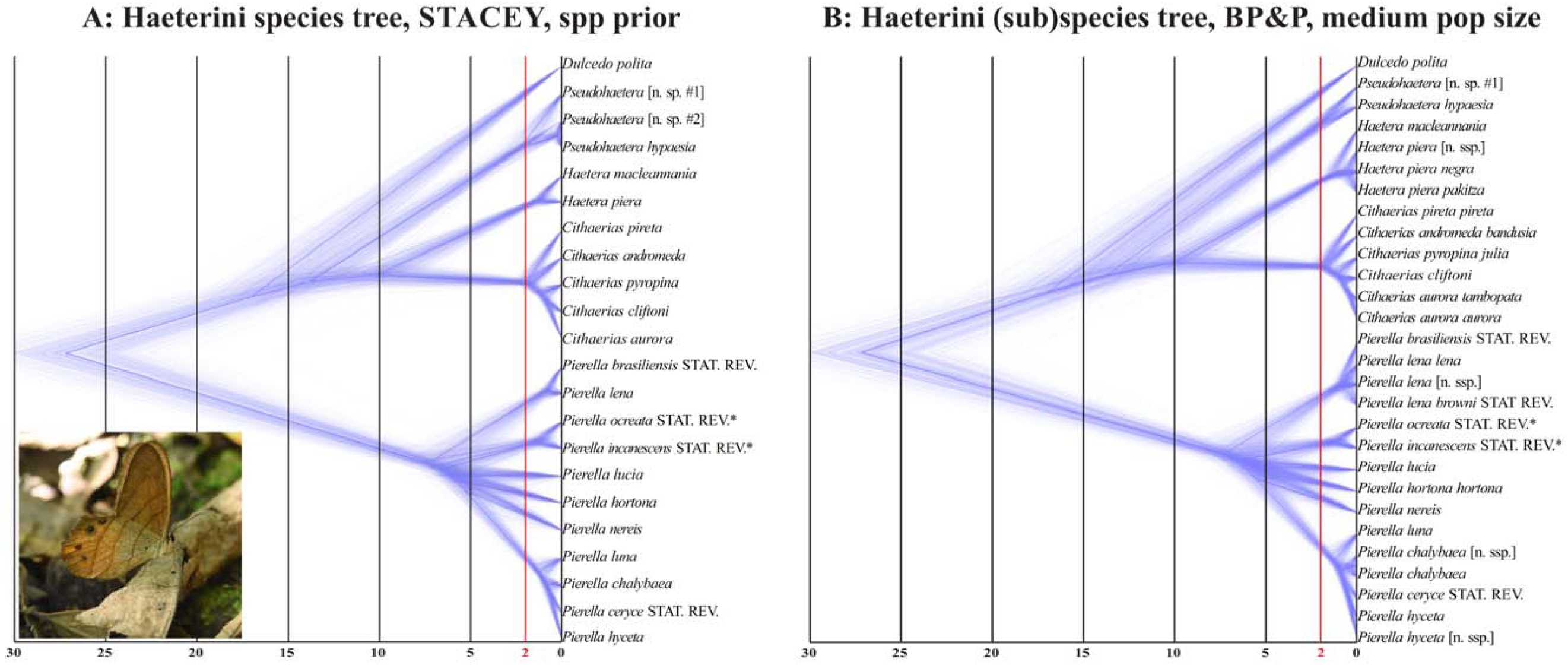
Time-calibrated phylogenetic hypotheses of models that best approximate species and subspecies in Haeterini. A) “Cloudogram” of the best-fit species delimitation model based on Bayes factors, STACEY under prior accounting for taxonomic species. B) “Cloudogram” of the delimitation model that best approximated described subspecies, BP&P under prior for medium ancestral population size. Time axis in both panels is scaled to million years. *The species status of *Pierella helvina ocreata* and *P. helvina incanescens* may change with the inclusion of *P. helvina helvina*, but it is likely that *P. helvina ocreata* and *P. helvina incanescens* are separate species. Inset butterfly: *Pierella hyceta hyceta*; Peru, Pasco, Cañón de Huancabamba, 1200 masl, 29.vii.2017. Photo: Markéta Aubrechtová.

## RESULTS

### Data compilation and potential phylogenetic biases

The single-gene and concatenated multi-locus tree topologies were congruent and showed no evident signature of cross-contamination (Fig. S1). The inferred inter-generic relationships were robust as indicated by high posterior probabilities (PP = 1.0 for the concatenated multi-locus tree) and the posterior MSC trees (Figs 2, 3). All clearwing Haeterini form a monophyletic group sister to the Haeterini butterflies having full scale-cover on wings. Therefore, in terms of phylogenetic branching pattern, the genus *Pierella* diverged early in the evolution of Haeterini, followed by the monotypic genus *Dulcedo*, the genus *Pseudohaetera*, and the divergence between the genera *Haetera* and *Cithaerias*. Mixed node support for inter-specific relationships were recovered in single-gene and in the concatenated multi-locus datasets, with PP ranging from ~0.6 to 1.0. The six loci chosen for this study have been previously utilized in butterfly species-level systematics, thus, we expected these loci to be phylogenetically informative (Table 2). Instead, we recovered low node support among certain Haeterini species (Fig. S1), such as those that rapidly diverged early in the evolution of *Pierella*, or in the recent radiation (< 2.5 Ma) of the genera *Pierella* and *Cithaerias*.

### Species delimitation

Regardless of which prior was used to take into account taxonomic knowledge for the collapseWeight parameter (ω), STACEY converged in similar MSCCs suggesting that either 22 (under parameter ω = 0.73) or 24 (under parameter ω = 0.59) lineages are the most adequate representation of species in our dataset. Indeed the pairwise similarity matrices generated for the two analyses are highly congruent, though the pairwise posterior probabilities seem to decrease in the STACEY analysis using ω = 0.59 (Fig. S2). On average, the posterior probability that two or more specimens inferred by STACEY under parameter ω = 0.73 belong to a single MSCC ranged from 0.42 to 0.99, with a median of 0.85 and mode of 0.93. The lowest posterior probability (0.42) was for the MSCC encompassing *Pierella lena lena*. In contrast, the STACEY analysis that did not take into account any taxonomy information (parameter ω with Beta distribution [α = 2, β = 2]) did not group any specimen in the dataset into MSCCs. The MSCCs inferred by BP&P showed high sensitivity to prior distributions for the parameter θ. The scenario with small ancestral population size, i.e. θ = IG[3, 0.02], suggested that nearly every specimen in the dataset represent a single divergent lineage. This extreme scenario suggesting 55 MSCCs was considered in the hypothesis testing exercise using Bayes factors, even though it significantly departs from current taxonomic understanding of Haeterini. The most likely numbers of MSCCs recovered by the other two BP&P analyses were 21 (when θ = IG[3, 0.2]; prior mean of 0.1) or 28 (when θ = IG[3, 0.1]; prior mean of 0.05). The two independent runs for each analysis converged in the same MSCCs, and the posterior probabilities of the delimited species for the analysis under θ = IG[3, 0.2] ranged from 0.44 to 1.00, with a median of 0.96 and mode of 0.99. The two lowest posterior probabilities (0.44) were for the MSCCs encompassing *Pierella lena lena* and for *Pierella hyceta hyceta*. Overall, the large impact of the prior distribution of the parameter θ in BP&P, resulting in 21, 28 or 55 MSCCs, suggests that more data, both genetic and taxon sampling, is needed to strongly inform the molecular species delimitation. However, the resulting most-probable number of MSCCs favored under the two scenarios for large and medium ancestral population sizes are congruent with the taxonomic knowledge of the group, as well as with the species delimitation exercises carried out in STACEY (Fig. 2).

The eight species delimitation hypotheses ranged from 18 to 63 species (or MSC clusters) (Fig. 2). The species hypotheses recovered by STACEY using ω = 0.73 (22 MSCCs) and BP&P using θ = IG[3, 0.2] (21 MSCCs) were strongly supported compared to the remaining six species delimitation hypotheses (*ln*Bf >> 10). The STACEY analysis recovering 22 MSCCs is supported, but not conclusively (*ln*Bf = 9.69), over the BP&P analysis recovering 21 MSCCs (Table 3). The differences between these two species hypotheses are: 1) the delimitation of two lineages of *Pseudohaetera hypaesia*, one from central Peru (Chanchamayo Valley) and the other from southeastern Ecuador (Morona Santiago and Zamora Chinchipe Provinces); BP&P suggested a single MSCC for these two lineages whereas STACEY suggested two separate MSCCs, 2) split of the subspecies *Pierella helvina ocreata* and *P. helvina incanescens* into two separate species only by STACEY, and 3) split of *Pierella chalybaea* from Ecuador (Sucumbios) from remaining *P. chalybaea* only by BP&P. On the other hand, the BP&P analyses under the medium ancestral population size (θ = IG[3, 0.1]) recovered most of the described subspecies in Haeterini, thus, we used this delimitation as a proxy for recognizing taxonomic subspecies in the framework of the MSC despite it not being supported by the Bayes factor model comparison (Fig. 2).

**Table 3:** Marginal-likelihood calculations using path sampling and Bayes factor model testing. Eight competing, non-nested species delimitation models were compared, and the STACEY analysis under prior accounting for the number of taxonomic species (*spp prior*) had the highest marginal likelihood estimate. Bayes factors (*ln*BF) = 2–10 were considered to represent positive support, while *ln*Bf > 10 were considered as decisive support.

| Delimitation model | Marginal L (Path Sampling) | Delimited species | $\ln\text{BF}$ STACEY, *spp prior* |
| --- | --- | --- | --- |
| STACEY, *spp prior* | -43522.54 | 22 | 0 |
| BP&P, large pop size | -43532.23 | 21 | 9.69 |
| Taxonomic *spp* | -43537.35 | 18 | 14.81 |
| STACEY, *sspp prior* | -43674.24 | 24 | 151.70 |
| BP&P, medium pop size | -43689.95 | 28 | 167.41 |
| Taxonomic *sspp* | -43701.81 | 26 | 179.27 |
| STACEY, no prior | -43719.44 | 63 | 196.90 |
| BP&P, small pop size | -43737.40 | 55 | 214.86 |

### Absolute divergence times

We show in Figure 3 two time-calibrated species trees, one approximating species in Haeterini (i.e. STACEY’s clusters under ω = 0.73; which had the highest marginal likelihood estimate among the eight species delimitation hypotheses) and the second approximating subspecies in Haeterini (BP&P’s clusters under θ = IG[3, 0.1]; which recovered most of the described taxonomic subspecies). The remaining six species trees can be found in the Supporting Information (Fig. S3). Species tree topologies and branch lengths in absolute time remained virtually the same regardless of species delimitation hypothesis. The median crown age of extant Haeterini was estimated at 27.2 Ma (95% HPD: 21.8 to 32.7 Ma; entire posterior range: 15.4 to 40.5 Ma). The rapid early radiation of *Pierella* happened in the late Miocene, at about 7.22 Ma (95% HPD: 5.68 to 9.03 Ma; entire posterior range: 4.97 to 11.4 Ma), whereas the recent radiation of *Pierella* and *Cithaerias* occurred in the Pleistocene, at about 1 to 2 Ma (95% HPD: 0.81 to 2.58 Ma; entire posterior ranges for the recent radiation of *Pierella*: 0.57 to 2.83 Ma and for *Cithaerias*: 1.19 to 3.25 Ma). We estimate that these two diversification events gave rise to about 70% of extant Haeterini species.

## DISCUSSION

Regardless of the species concept advocated by different researchers, traditional taxonomy and coalescent-based approaches can act in synergy to infer statistically robust species hypotheses. Importantly, the multispecies coalescent (MSC) model and the Bayesian framework to delimit MSC clusters (MSCCs) allow the quantification and testing of species boundaries informed by taxonomic knowledge. Furthermore, divergent taxonomic opinion among authors working on particular species groups has been statistically weighed by the approach followed in this study. This is important as a first step towards reliable standardization of taxonomy and higher-level systematics based on models that take into account the process of speciation (e.g. incomplete lineage sorting) and not just arbitrary genetic distances as in threshold-based approaches (e.g. based on COI barcoding). Here, we have not used any threshold-based method because it would not contribute to the described probabilistic framework as these methods utilize single-locus datasets and neglect tree topology uncertainty. However, future research can further address comparisons between threshold-based approaches (e.g., single-threshold, multi-thresholds, multi-rate processes) and the multispecies coalescent model.

More realistic models of speciation that include, for example, inter-specific gene flow might be more accurate at estimating species histories (Müller *et al.*, 2018). For example, in Europe alone, around 16% of butterfly species are known to hybridize (Descimon & Mallet, 2009), which is in line with the idea of speciation as a continuum (De Queiroz, 2007) and that inter-specific gene flow in animals might not be uncommon (Mallet *et al.*, 2007). Another source of information for taxonomic conclusions is the development of new methods that jointly model phenotypic traits and genetic data (Solís-Lemus *et al.*, 2015). Unfortunately, phenotypic and life history data, such as larval host plants and habitat association, that could be used in species delimitation is scarce for Haeterini butterflies. Taken together, these methodological advances and the generation of biological data from various sources suggest that in the near future coalescent-based approaches based on multi-locus data may be able to overcome many current shortcomings in delimiting species.

### Limitations and strengths

The dataset and approach that we have followed here are not exempt of limitations. First, the amount of missing information, including taxonomic sampling and gapped molecular dataset, may have reduced the power of our analyses. However, it has been noted previously that the MSC might still be accurate with sampling schemes including fewer than five individuals per lineage, as long as multi-locus datasets (in our case 6 unlinked loci) are utilized (Zhang *et al.*, 2011). Although the impact of missing data in the MSC framework needs to be further examined, a recent simulation study suggested that coalescent-based species tree inference might be highly accurate even with severely gapped multi-locus datasets (Nute *et al.*, 2018). Second, inter-specific gene flow may impact current implementations of the MSC by obscuring lineages divergence (Luo *et al.*, 2018). Nevertheless, unless there is a high level of gene flow and hybridization, the MSC-based delimitation and species tree inference should be robust (Eckert & Carstens, 2008; Zhang *et al.*, 2011). However, we note that this issue needs to be further studied in the light of more data and using recently-developed approaches, such as the isolation-with-migration model (Müller *et al.*, 2018). Third, the impact of priors in MSC-based species delimitation might be high when the molecular dataset does not hold sufficient signal to converge on the same MSCCs (Leaché & Fujita, 2010; Jones *et al.*, 2015). Our results showed that the choice of priors heavily influenced the number of MSCCs estimated by MSC-based methods. However, our priors rely on existing taxonomic knowledge and thus we have reduced the clustering space of 63 specimens based on actual evidence coming from previous morphology-based studies. The number of most likely MSCCs inferred here (from 21 to 28 clusters) remains highly congruent with the morphological diversity encountered in this group of butterflies.

The strengths of our study rely on three key aspects. First, we have not used any a priori taxonomic assignment of individuals to species, nor any guide species tree to delimit species. This allowed us to include both tree topology and taxonomic assignment uncertainties explicitly into the models, avoiding potential biases that may have precluded accurate estimation of speciation probabilities (Leaché & Fujita, 2010). However, we note that the posterior probabilities of certain species delimitations, such as *Pierella lena lena*, remain low, which may be explained by non-sufficient signal of our molecular dataset. Adding more genetic data may increase the posterior probabilities while holding the same delimitations, given that different approaches converged in the same species hypotheses regardless of the low posterior probabilities (e.g. *Pierella lena lena* was recovered as most probable by both STACEY and BP&P, Fig. 2). However, this needs to be confirmed by future genome-wide studies on Haeterini. Second, taxonomic knowledge has been formally taken into account in our probabilistic scenarios. Previous studies have put forward the statistical comparison (e.g. via Bayes factors) of alternate species delimitation models, but these mostly evaluated different individual reassignments based only on node collapsing of sister lineages in a phylogeny (Grummer *et al.*, 2014; Yu *et al.*, 2017). Our pipeline, on the contrary, has aided the exploration of alternate MSCCs that are congruent as well with other sources of information in butterfly taxonomy, such as morphology and geography. Third, the MSCCs found in this study are not nested, meaning that specimens could be re-assigned to any combination of clusterings. The use of Bayes factors as a selection tool is appropriate because of its flexibility in testing non-nested models (Leaché *et al.*, 2014). We used path sampling under thermodynamic integration which has been shown to be highly accurate in testing non-nested species delimitation models (Grummer *et al.*, 2014).

### Performance of methods

Our approach focused on using the multispecies coalescent model, informed by taxonomic knowledge, to assign individuals to MSCCs based on different priors and methods implemented in STACEY and BP&P. This is arguably a less arbitrary approach to reduce the space of all possible clusterings for model testing using Bayes factors, compared to other approaches based solely in taxonomic expertise (Leaché *et al.*, 2014) or in multi-locus networks (Grummer *et al.*, 2014) and population assignments (Olave *et al.*, 2014). Furthermore, the approach outlined here is flexible, simple and fast because it avoids preliminary estimation of guide trees and *a priori* taxonomic assignments of individuals. Taxonomic knowledge modeled as prior distribution has the potential to ameliorate potential biases (e.g. population structure raised to species), which have been observed when relying solely on genetic information and guide trees (Olave et al., 2014). As Sukumaran & Knowles (2017) argue, the only way that the multispecies coalescent can be used as a reliable species delimitation approach is to explicitly incorporate other sources of information as *a priori* hypotheses. We showed that the number of inferred MSCCs heavily depends on priors, and indeed, under the scenario of medium ancestral population sizes (θ = IG[3, 0.1]) BP&P recovered traditionally viewed infra-specific diversity, and perhaps structured populations. Note that subspecies in butterfly taxonomy are seen as populations with limited gene flow due to allopatric distributions (Lamas, 2008). Overall, we highlight the importance of taxonomically-informed molecular species delimitation and the use of Bayes factor model comparison.

Both MSC-based methods used in this study, STACEY and BP&P, were fast and simple in their implementation thanks to available guidelines and manuals (e.g. Jones *et al.*, 2015; Yang, 2015). The most supported BP&P’s delimitation followed a conservative prior distribution for the size of ancestral population size, that is θ = IG[3, 0.2] with a prior mean = 0.1, which was pointed out as the most appropriate prior mean (Leaché & Fujita, 2010). This species delimitation model is highly similar to the most supported STACEY model, except for three species hypotheses. In line with morphological differences, STACEY favored the split of *Pierella helvina incanescens* and *P. helvina ocreata*, two subspecies that were originally considered separate species but synonymized due to allopatric geographical distribution by Constantino (1995). On the other hand, BP&P favored the split of one *Pierella chalybaea* specimen (Ecuador, Sucumbíos) from other conspecific individuals, including others from Ecuador. STACEY did not favor such a split, which again is in line with the absence of any clear morphological difference in *P. chalybaea* from western Amazonia (Zacca *et al.*, 2016). Finally, STACEY delimited three separate species within *Pseudohatera hypaesia* while BP&P delimited only two. However, there are no clear wing coloration differences among populations of *P. hypaesia* throughout its range from Colombia to Bolivia (Gerardo Lamas, *pers. comm.*). The supported STACEY’s delimitation here thus points out that large genetic divergences exist in the otherwise similar-looking *P. hypaesia*, warranting a taxonomic revision of these montane butterflies from the tropical Andes.

### Taxonomic implications

The probabilistic framework applied in this study allows the statistical test of alternative species hypotheses in a taxonomic group that has likely evolved for nearly 27 million years. The most likely scenario among those tested here suggests that at least four divergent lineages should be elevated to species by current taxonomic standards. It is clear that a more comprehensive sampling and datasets, including morphological and molecular characters, are needed to robustly delineate species boundaries in Haeterini. Nonetheless, the four divergent lineages found in this study corroborate morphological differences that have been previously acknowledged in these groups but that taxonomists conservatively considered conspecific variations (subspecies) mainly because of allopatric distributions, a historical practice in butterfly taxonomy (Lamas, 2008). In almost all MSC-based analyses, the following taxa were considered independent evolutionary lineages: i) *Pierella hyceta hyceta* and *Pierella hyceta ceryce*, ii) *Pierella lena lena* and *Pierella lena brasiliensis*, iii) *Pierella helvina incanescens* and *Pierella helvina ocreata*, iv) two divergent sympatric populations in central Peru (Chanchamayo Valley) of *Pseudohaetera hypaesia*. According to our results, these eight lineages are likely full species (a taxonomic revision in preparation is addressing this).

Zacca *et al.* (2016) split *Pierella lamia* into seven species mainly based on allopatric geographical distributions, and also on genitalia and androconial patches on male wings (despite their similar wing coloration). We did not find support for the species status of *Pierella keithbrowni* from southeastern Brazil proposed by these authors. In the case of *P. keithbrowni*, the main diagnostic characters were the androconial patch shape, and ductus bursae in female genitalia longer than in *P. nice*, albeit male genitalia in *Pierella lamia* complex are not differentiated. In all MSC-based analyses, *P. keithbrowni* is not genetically different from other populations in central Brazil and Ecuador. Our results therefore suggest that differences in the aforementioned morphological characters may represent variation within a single evolutionary lineage, and thus their usage needs to be complemented with other lines of evidence. Note that androconial patch morphology has been widely used to diagnose butterfly species boundaries in other satyrine groups (e.g. Núñez Aguila *et al.*, 2013; Penz *et al.*, 2017). Our results question the reliability of this character system only in Haeterini lineages, thus, further research in other butterfly groups can test the generality of our conclusions that different character systems favor narrower or broader delimitations in butterflies. However, as noted previously, granting species status to butterfly populations primarily based on geographical distribution (allopatric populations) might be unjustified if other evidence fails to recognize clear divergence (Descimon & Mallet, 2009). Therefore, we suggest that *P. keithbrowni* should be synonymized with *P. chalybaea* (work in preparation will address this).

## CONCLUSIONS

Here we provide evidence that for the butterfly tribe Haeterini, the multispecies coalescent model generally recognizes traditionally viewed butterfly subspecies and species, with some exceptions linked to the use of different morphological character systems. By using a probabilistic framework, we show that divergent taxonomic opinion (concepts) were used by different authors, including butterfly species that were over-split (*lamia* complex), lumped (at least 6 subspecies raised to species here), or misclassified (e.g. *Pierella lena browni* was previously synonymized with *P. lena brasiliensis* (Lamas, 2004), despite *browni* being evolutionarily more closely related to *P. lena lena* than to *brasiliensis*). Androconial patch morphology is commonly used as informative character to diagnose species, but we show that at least in *Pierella* it alone may not be well-suited to distinguish infra- and inter-specific diversity. Furthermore, taxonomic knowledge informed as priors in MSC-based species delimitation using genetic data is a robust approach to reduce the clustering space in model testing.

The low node support recovered among certain Haeterini species may be attributed to incomplete lineage sorting due, for example, to ancient rapid radiation as in the crown node of *Pierella* or to recent speciation as in *Pierella* and *Cithaerias*. Haeterini butterflies evolved for nearly 27 million years but most extant species (*ca*. 80%) likely diverged rather recently—within the past 2 million years. Future macroevolutionary studies using the revised species boundaries of Haeterini might address this puzzling diversification history, whether it was the result of high tropical species turnover over millions of years—with constant and high extinction and speciation rates, in line with Antonelli *et al.* (2015) —or a Pleistocene major burst in diversification, in line with the Quaternary diversification model, as characterized via simulated data (Matos-Maraví, 2016).

## SUPPLEMENTARY MATERIAL

**Figure S1:** Consensus trees based on single-gene and concatenated loci datasets (pages 2–9).

**Figure S2:** Pairwise similarity matrices based on delimitation analyses with STACEY (pages 10– 14).

**Figure S3:** Time-calibrated Maximum Clade Credibility species trees of the eight species delimitation hypotheses (pages 15–24).

DNA alignments and phylogenies of the single-gene and concatenated datasets were archived in TreeBASE (ID: 23439). Accession URL:

http://purl.org/phylo/treebase/phylows/study/TB2:S23439.

Input files for species delimitation in BP&P and STACEY, and time-calibrated phylogenies: FigShare, https://doi.org/10.6084/m9.figshare.7637021.v1.

## ACKNOWLEDGEMENTS

We are grateful to Keith R. Willmott, André V.L. Freitas, Carlos Peña, Yves Basset and Phil DeVries for contributing specimens for this study, and to Yves Basset, Diana Silva, Gerardo Lamas and SERFOR (Ministerio de Agricultura, Peru; No Autorización 223-2017-SERFOR/DGGSPFFS) for assistance with research permits. We thank Graham R. Jones for assistance in running and interpreting results from STACEY. We thank Thomas Simonsen and two anonymous reviewers for constructive comments on a previous version of the manuscript.

