## SUPPLEMENTARY MATERIAL for "Species limits in butterflies (Lepidoptera: Nymphalidae): Reconciling classical taxonomy with the multispecies coalescent"

Figure S1: Single gene and concatenated loci trees. Each tree represents the 50% majority-rule consensus of 7,500 posterior sampled trees inferred in MrBayes. Posterior probabilities on each node are represented by colored circles following the figures' legend. A: CAD-gene tree; B: COI-gene tree; C: EF1α-gene tree; D: GAPDH-gene tree; E: RpS5-gene tree; F: *wingless*-gene tree; G: 6-loci concatenated tree.

A: CAD-gene tree

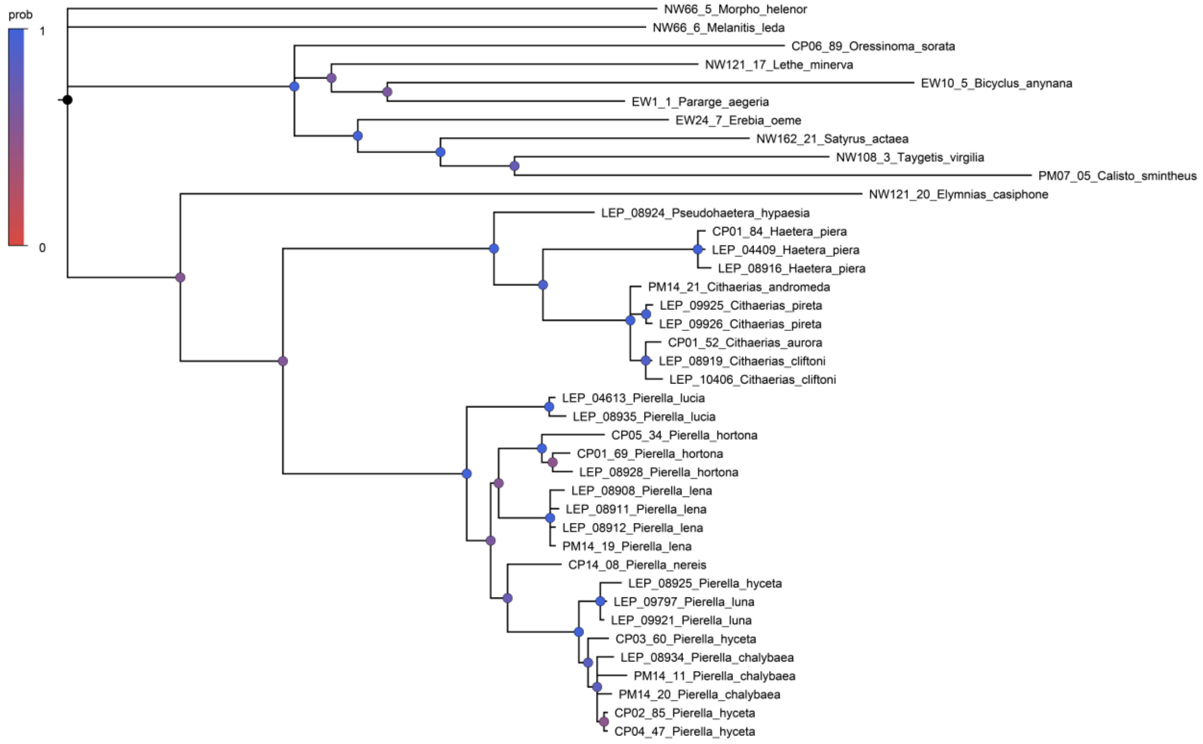

B: COI-gene tree

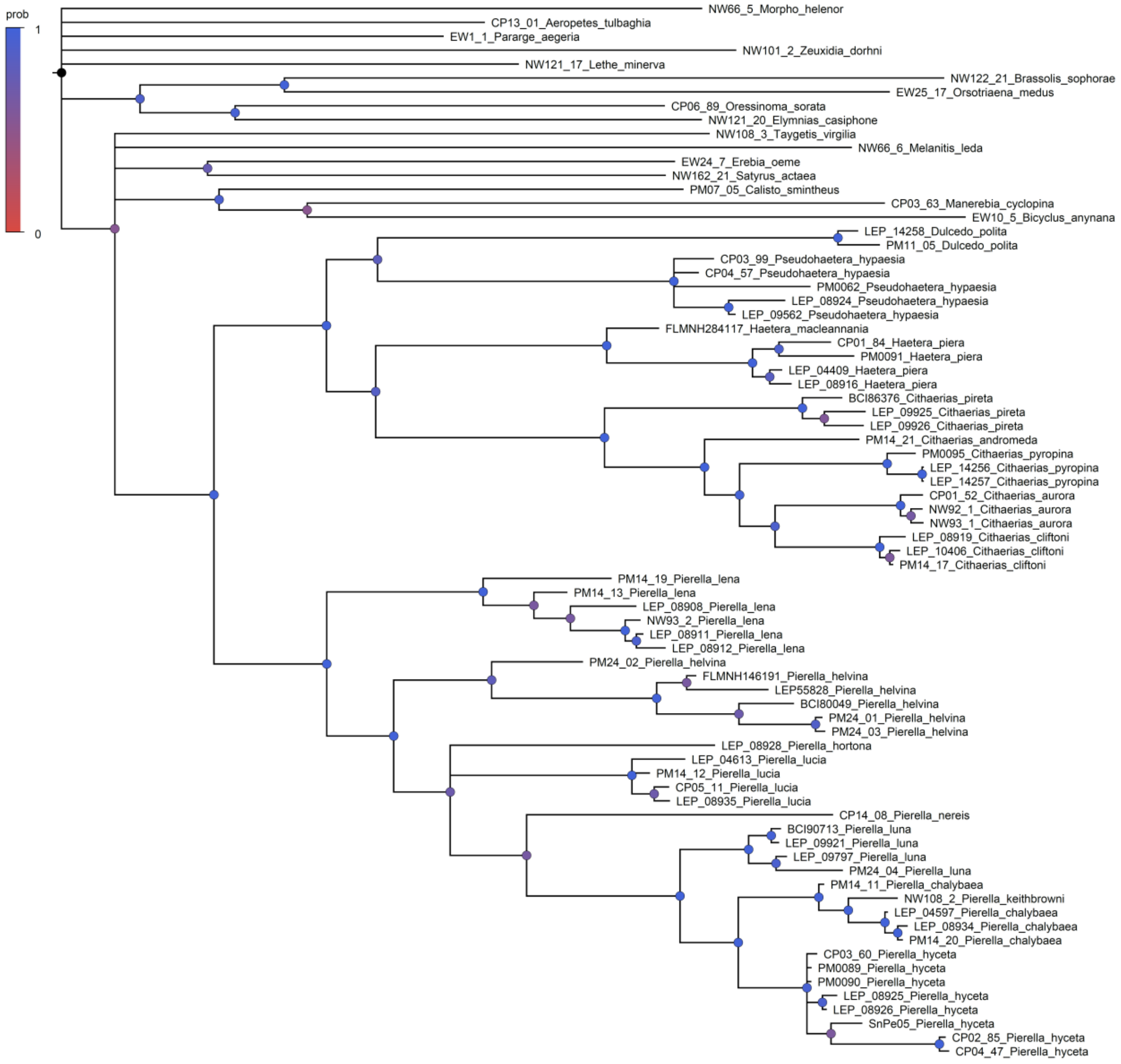

C: EF1α-gene tree

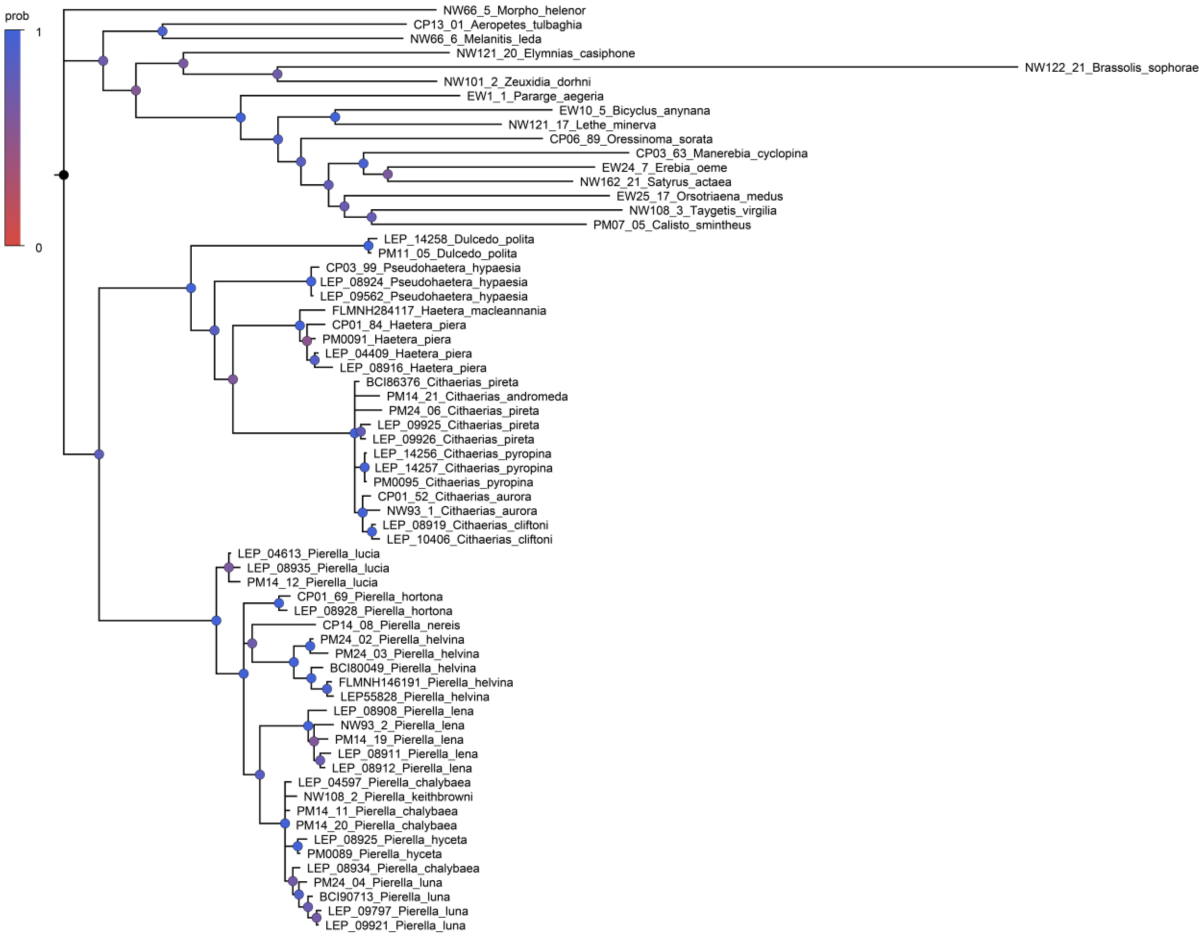

D: GAPDH-gene tree

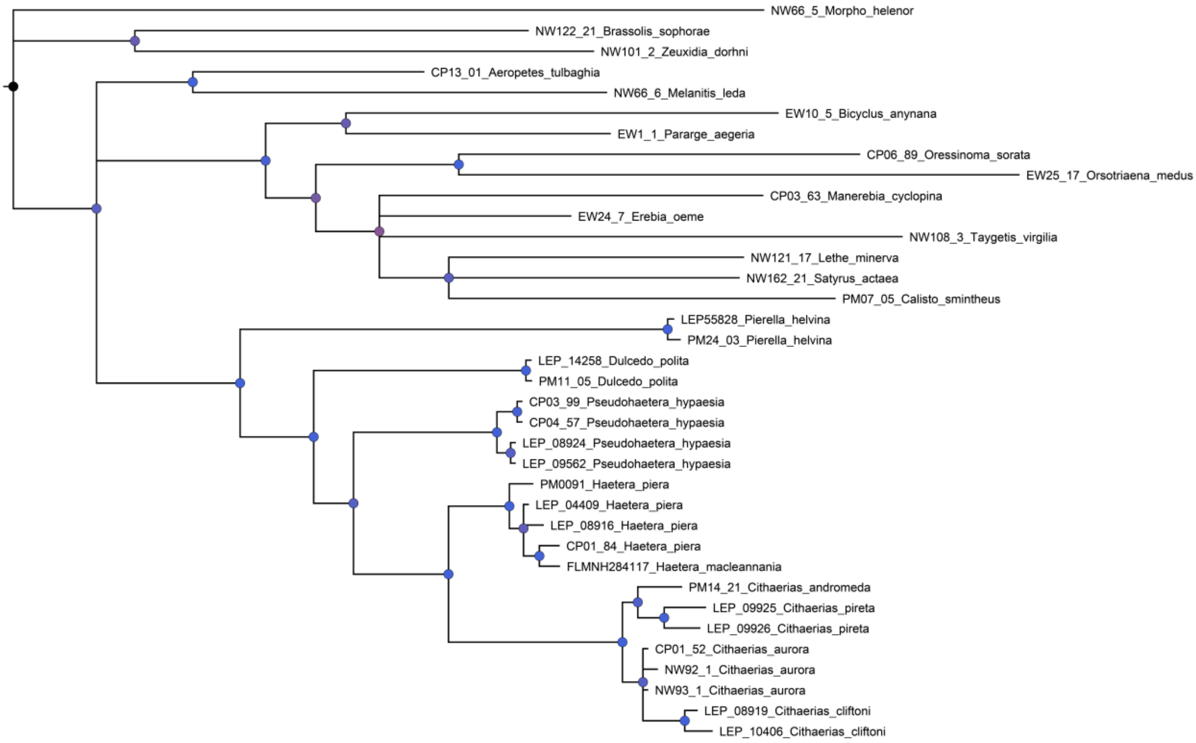

E: RpS5-gene tree

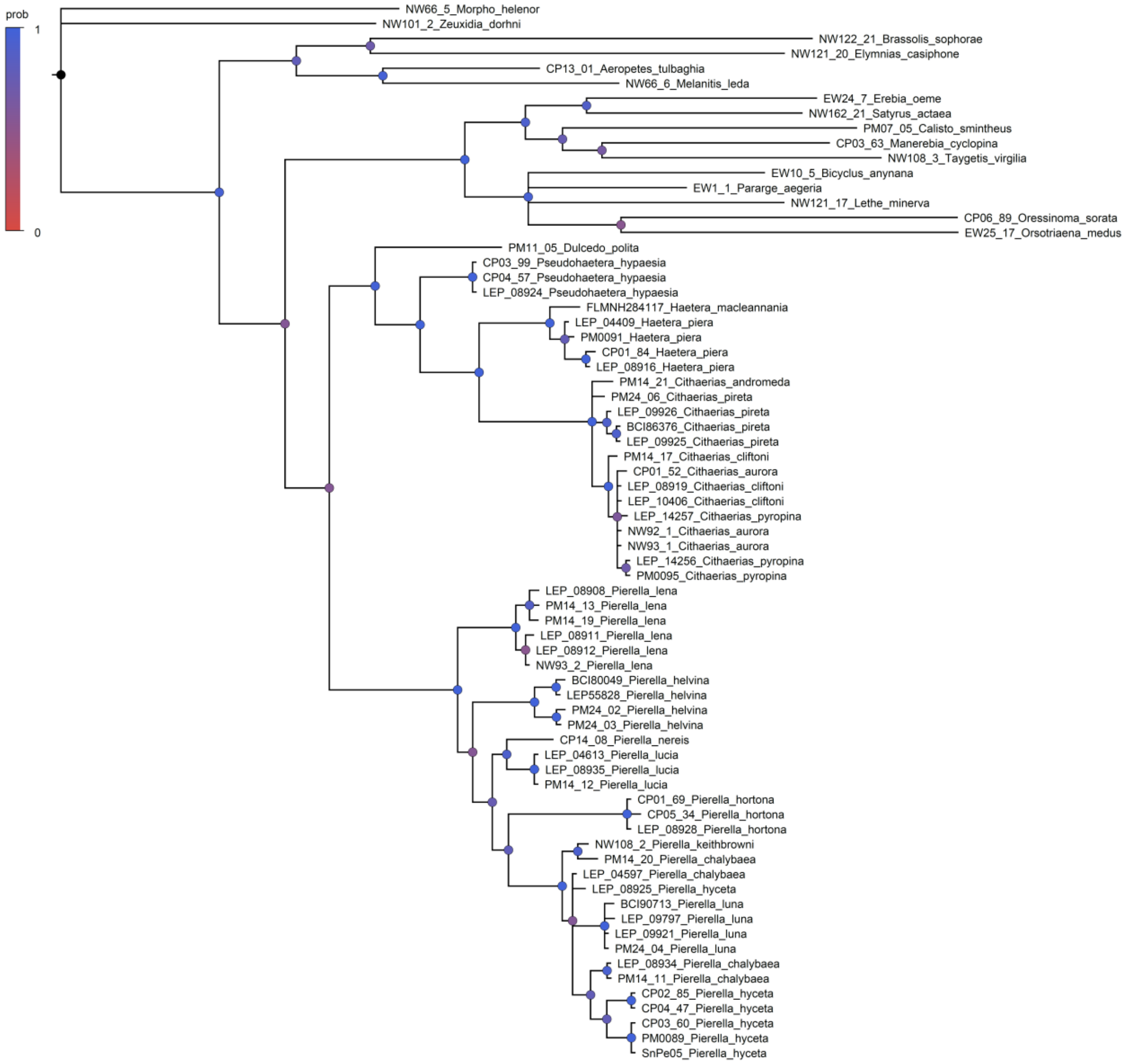

F: *wingless*-gene tree

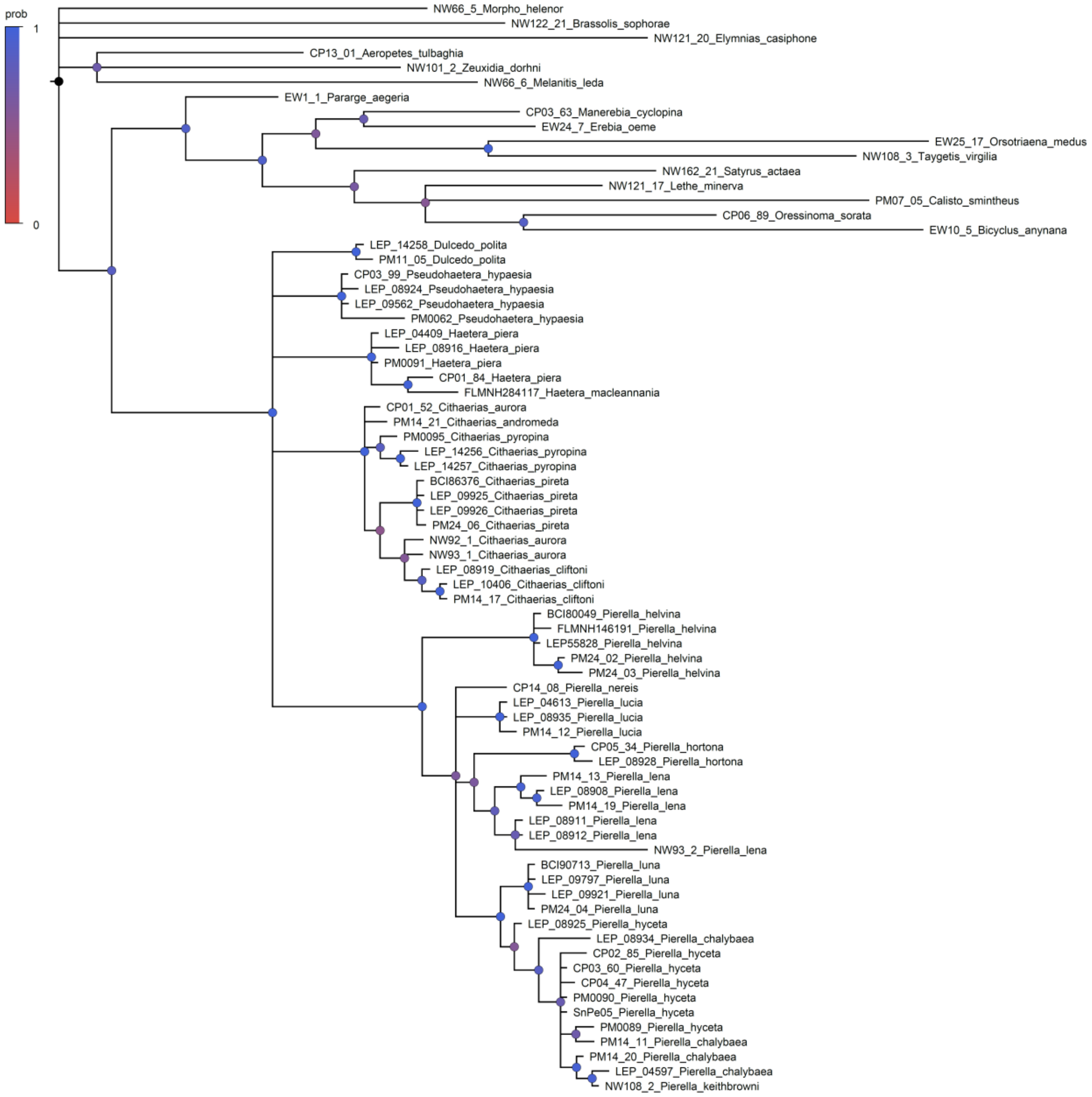

G: 6-loci concatenated tree

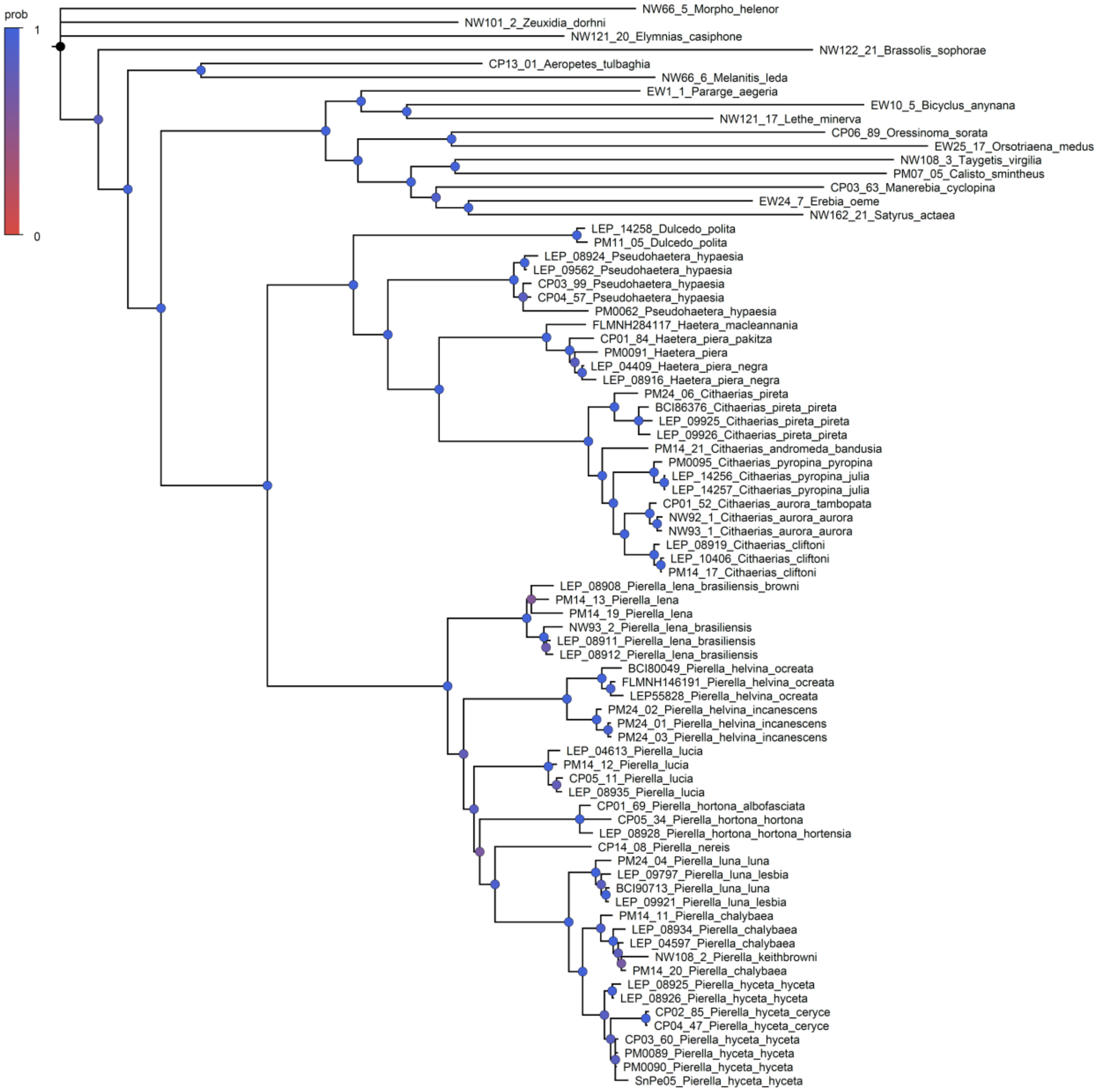

Figure S2: Pairwise similarity matrix among all 63 Haeterini sampled specimens. "Cloudograms" and species delimitations are the same as in Figure 2. Colored cells by similarity value follow the figure's legend. A: Posterior values from STACEY's delimitation model under $\omega = 0.73$ (spp prior following the number of taxonomic species); B: Posterior values from STACEY's delimitation model under $\omega = 0.59$ (sspp prior following the number of taxonomic subspecies elevated to species).

A: Posterior values from STACEY's delimitation model under ω = 0.73 (spp prior)

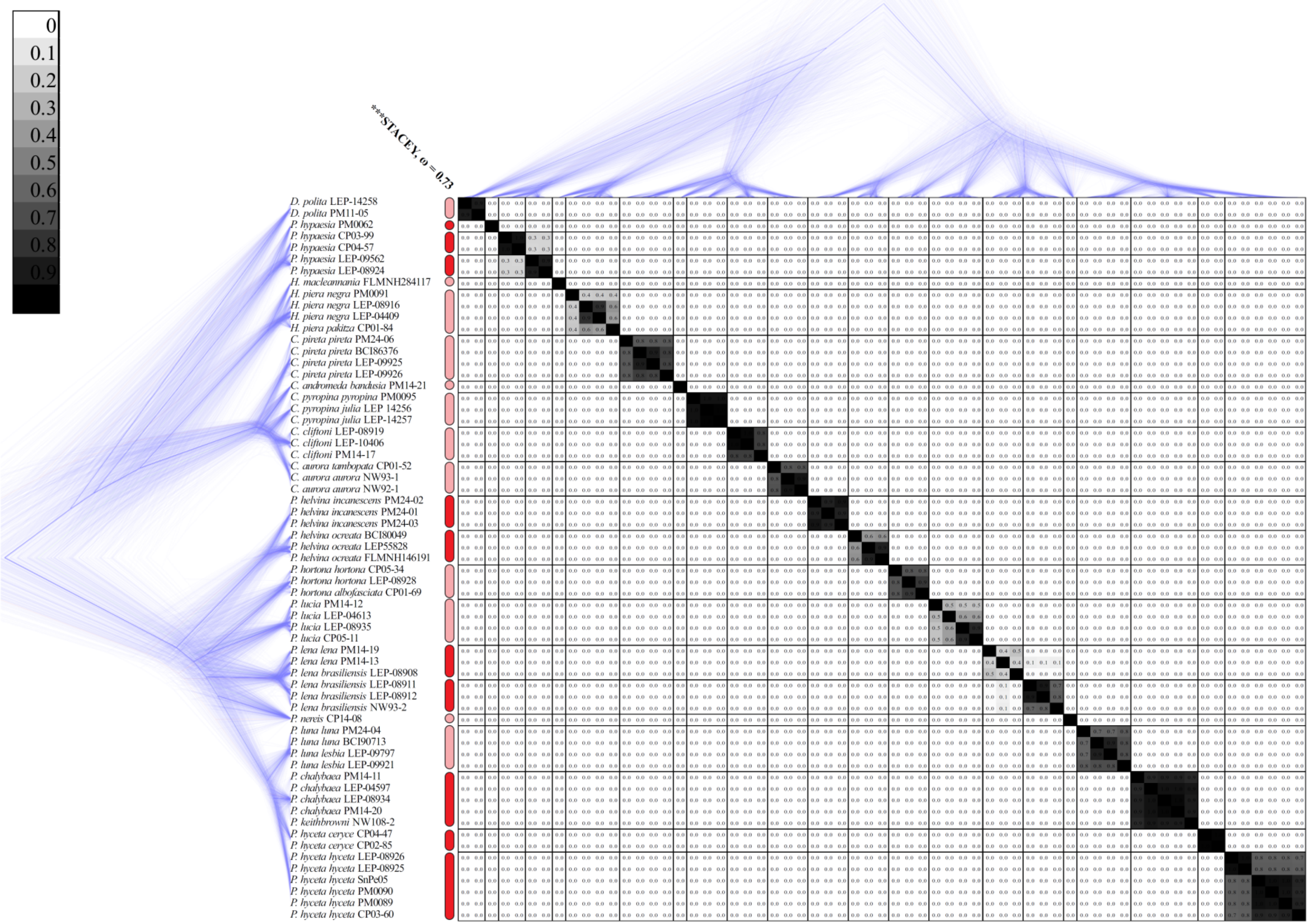
0
0.1
0.2
0.3
0.4
0.5
0.6
0.7
0.8
0.9
***STACEY, ω = 0.73
D. polita LEP-14258
D. polita PM11-05
P. hypaesia PM0062
P. hypaesia CP03-99
P. hypaesia CP04-57
P. hypaesia LEP-09562
P. hypaesia LEP-08924
H. macleannania FLMNH284117
H. piera negra PM0091
H. piera negra LEP-08916
H. piera negra LEP-04409
H. piera pakitza CP01-84
C. pireta pireta PM24-06
C. pireta pireta BCI86376
C. pireta pireta LEP-09925
C. pireta pireta LEP-09926
C. andromeda bandusia PM14-21
C. pyropina pyropina PM0095
C. pyropina julia LEP 14256
C. pyropina julia LEP-14257
C. cliftoni LEP-08919
C. cliftoni LEP-10406
C. cliftoni PM14-17
C. aurora tambopata CP01-52
C. aurora aurora NW93-1
C. aurora aurora NW92-1
P. helvina incanescens PM24-02
P. helvina incanescens PM24-01
P. helvina incanescens PM24-03
P. helvina ocreata BCI80049
P. helvina ocreata LEP55828
P. helvina ocreata FLMNH146191
P. hortona hortona CP05-34
P. hortona hortona LEP-08928
P. hortona albofasciata CP01-69
P. lucia PM14-12
P. lucia LEP-04613
P. lucia LEP-08935
P. lucia CP05-11
P. lena lena PM14-19
P. lena lena PM14-13
P. lena brasiliensis LEP-08908
P. lena brasiliensis LEP-08911
P. lena brasiliensis LEP-08912
P. lena brasiliensis NW93-2
P. nereis CP14-08
P. luna luna PM24-04
P. luna luna BCI90713
P. luna lesbia LEP-09797
P. luna lesbia LEP-09921
P. chalybaea PM14-11
P. chalybaea LEP-04597
P. chalybaea LEP-08934
P. chalybaea PM14-20
P. keithbrowni NW108-2
P. hyceta ceryce CP04-47
P. hyceta ceryce CP02-85
P. hyceta hyceta LEP-08926
P. hyceta hyceta LEP-08925
P. hyceta hyceta SnPe05
P. hyceta hyceta PM0090
P. hyceta hyceta PM0089
P. hyceta hyceta CP03-60

2

B: Posterior values from STACEY's delimitation model under $\omega = 0.59$ (sspp prior)

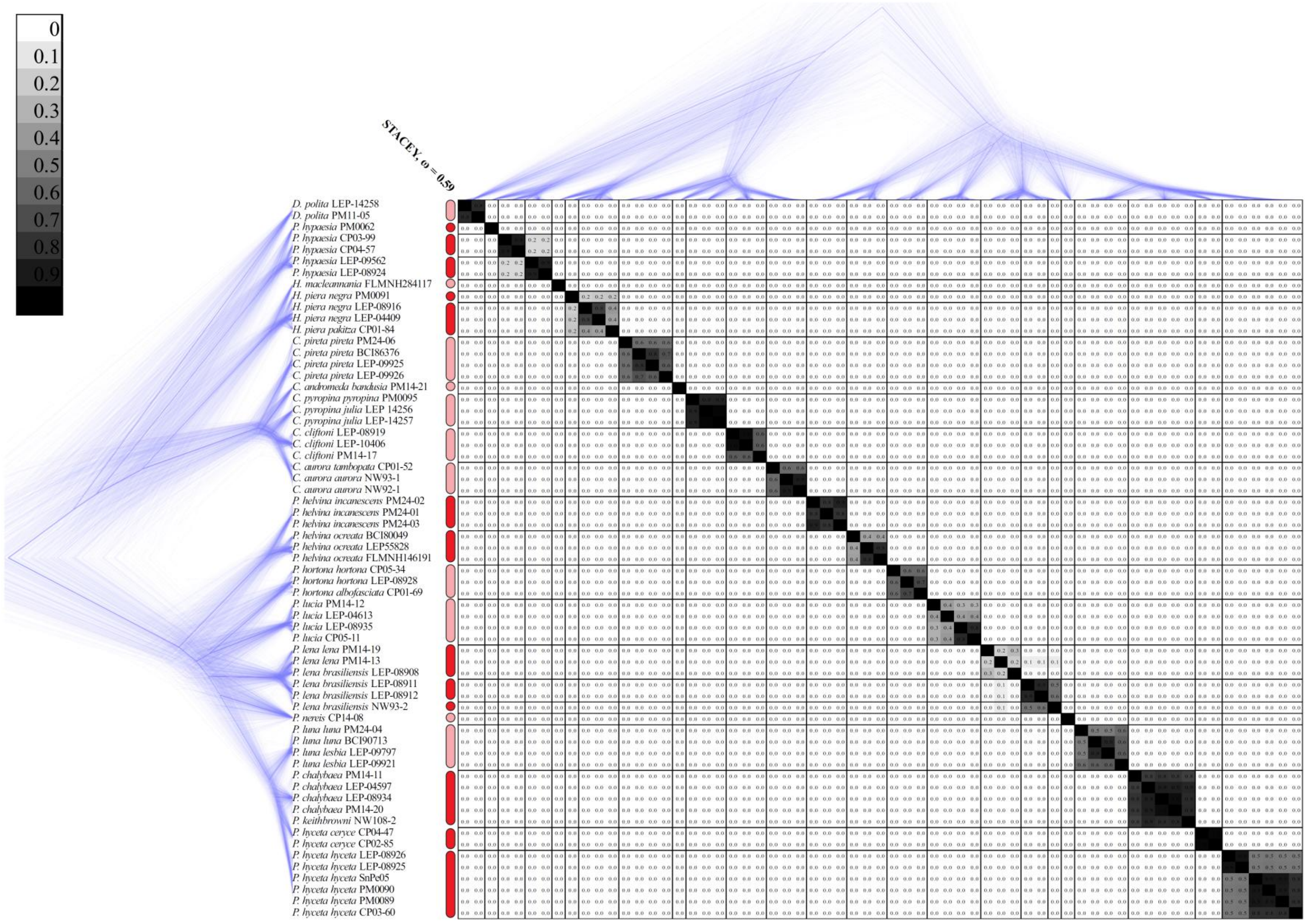
0
0.1
0.2
0.3
0.4
0.5
0.6
0.7
0.8
0.9
STACEY, ω = 0.59
D. polita LEP-14258
D. polita PM11-05
P. hypaesia PM0062
P. hypaesia CP03-99
P. hypaesia CP04-57
P. hypaesia LEP-09562
P. hypaesia LEP-08924
H. macleannania FLMNH284117
H. piera negra PM0091
H. piera negra LEP-08916
H. piera negra LEP-04409
H. piera pakitza CP01-84
C. pireta pireta PM24-06
C. pireta pireta BCI86376
C. pireta pireta LEP-09925
C. pireta pireta LEP-09926
C. andromeda bandusia PM14-21
C. pyropina pyropina PM0095
C. pyropina julia LEP 14256
C. pyropina julia LEP-14257
C. cliftoni LEP-08919
C. cliftoni LEP-10406
C. cliftoni PM14-17
C. aurora tambopata CP01-52
C. aurora aurora NW93-1
C. aurora aurora NW92-1
P. helvina incanescens PM24-02
P. helvina incanescens PM24-01
P. helvina incanescens PM24-03
P. helvina ocreata BCI80049
P. helvina ocreata LEP55828
P. helvina ocreata FLMNH146191
P. hortona hortona CP05-34
P. hortona hortona LEP-08928
P. hortona albofasciata CP01-69
P. lucia PM14-12
P. lucia LEP-04613
P. lucia LEP-08935
P. lucia CP05-11
P. lena lena PM14-19
P. lena lena PM14-13
P. lena brasiliensis LEP-08908
P. lena brasiliensis LEP-08911
P. lena brasiliensis LEP-08912
P. lena brasiliensis NW93-2
P. nereis CP14-08
P. luna luna PM24-04
P. luna luna BCI90713
P. luna lesbia LEP-09797
P. luna lesbia LEP-09921
P. chalybaea PM14-11
P. chalybaea LEP-04597
P. chalybaea LEP-08934
P. chalybaea PM14-20
P. keithbrowni NW108-2
P. hycetaceryce CP04-47
P. hyceta ceryce CP02-85
P. hyceta hyceta LEP-08926
P. hyceta hyceta LEP-08925
P. hyceta hyceta SnPe05
P. hyceta hyceta PM0090
P. hyceta hyceta PM0089
P. hyceta hyceta CP03-60

Figure S3: Time-calibrated Maximum Clade Credibility (MCC) species trees for eight species delimitation hypotheses. Each tree summarizes 7,500 posterior sampled trees inferred in starBEAST2, and specimens grouped following the eight delimitation models in Figure 2 in the main text. Posterior probabilities on each node are represented by colored circles following the figures' legend. 95% HPD (credibility interval) on each node is represented by blue horizontal bar. Time axis scaled to million years. A: Valid described species (taxonomic spp); B: Valid described subspecies (taxonomic sspp); C: STACEY's delimitation model under $\omega = 0.73$ (spp prior); D: STACEY's delimitation model under $\omega = 0.59$ (sspp prior); E: STACEY's delimitation under $\omega = \text{Beta}(2, 2)$ (no taxonomic prior); F: BP&P's delimitation model under $\theta = \text{IG}[3, 0.2]$ (large ancestral population size); G: BP&P's delimitation model under $\theta = \text{IG}[3, 0.1]$ (medium pop size); H: BP&P's delimitation model under $\theta = \text{IG}[3, 0.02]$ (small pop size).

A: Valid described species (taxonomic spp)

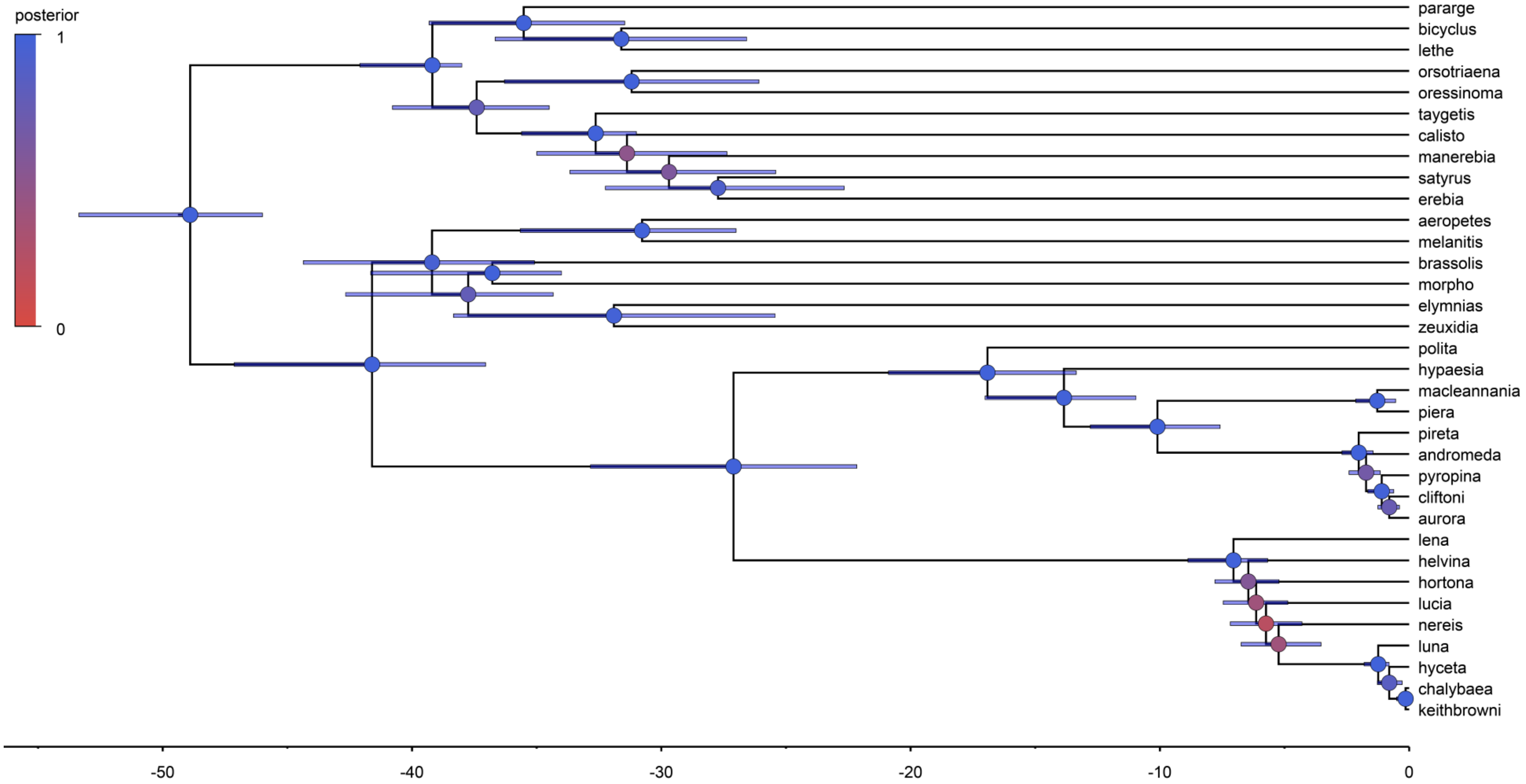

B: Valid described subspecies (taxonomic sspp)

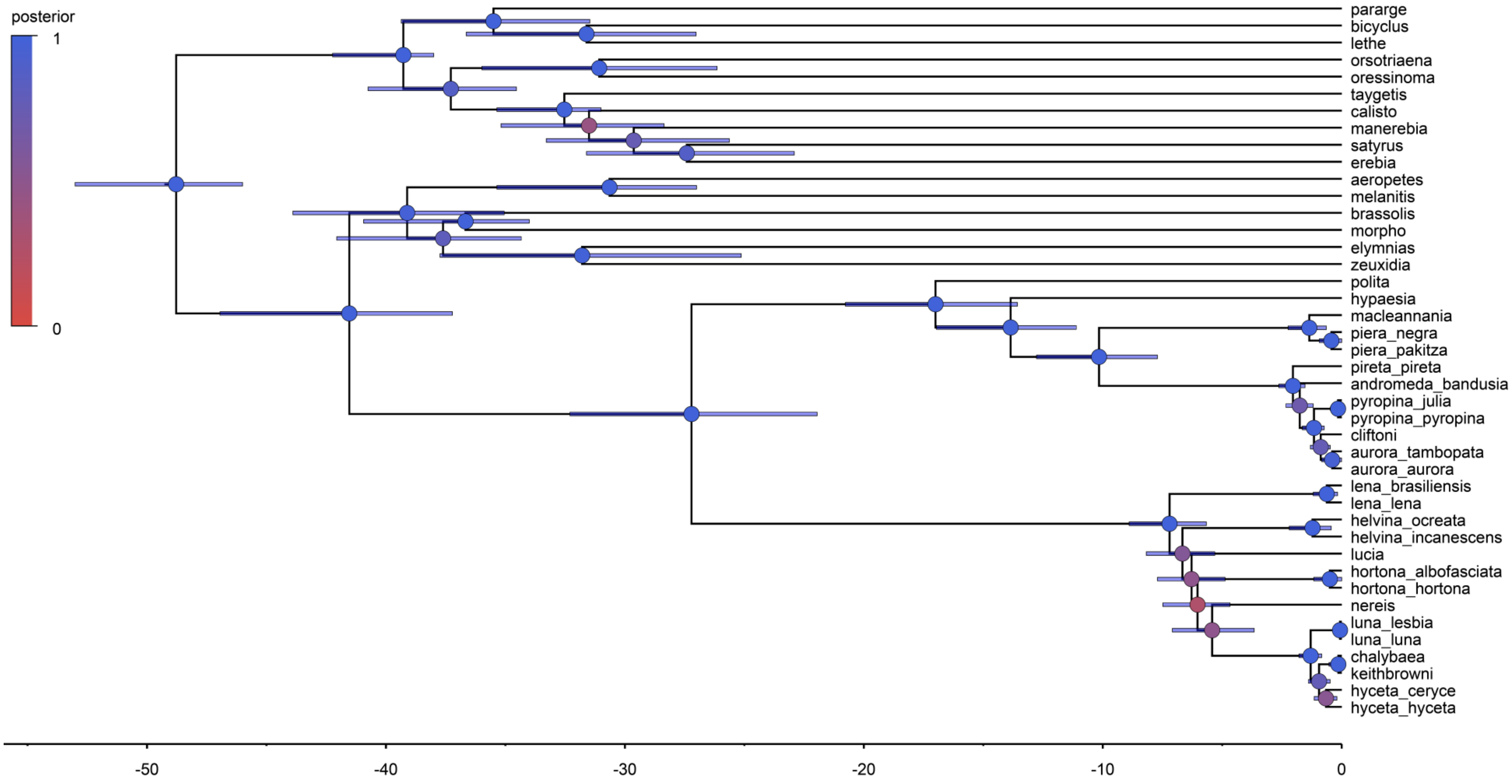

C: STACEY's delimitation model under $\omega = 0.73$ (spp prior)

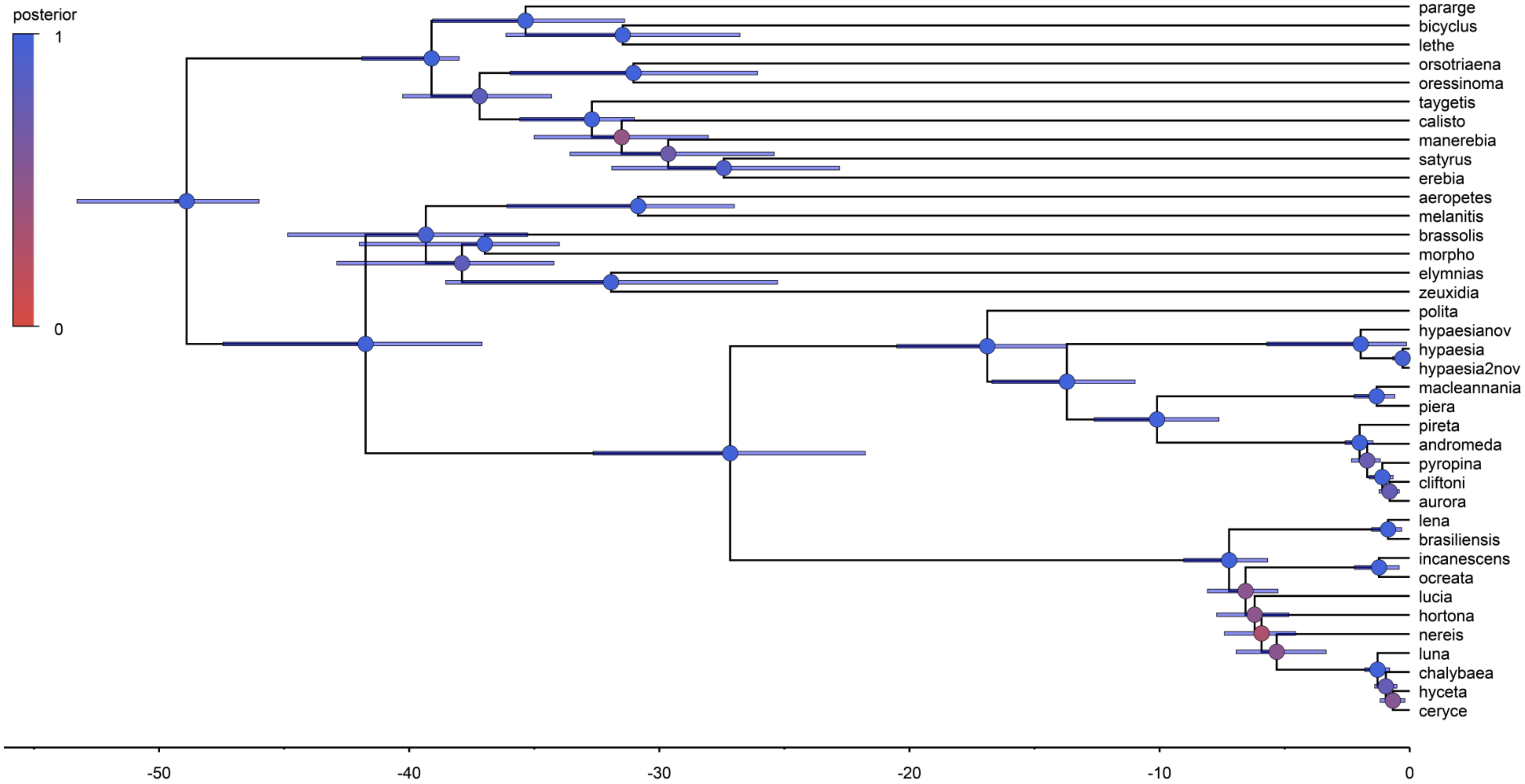

D: STACEY's delimitation model under ω = 0.59 (sspp prior)

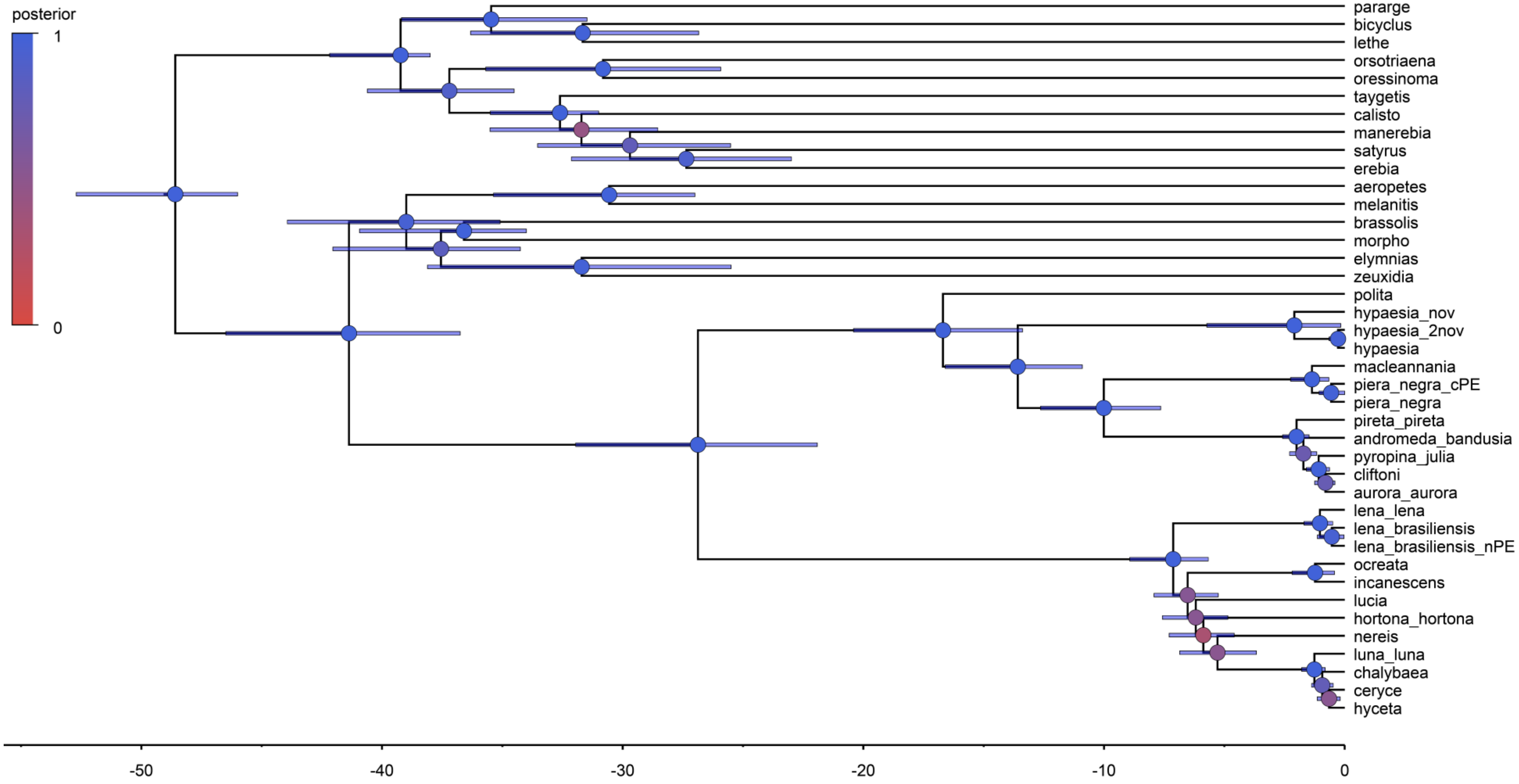

E: STACEY's delimitation model under ω = Beta(2, 2) (no prior)

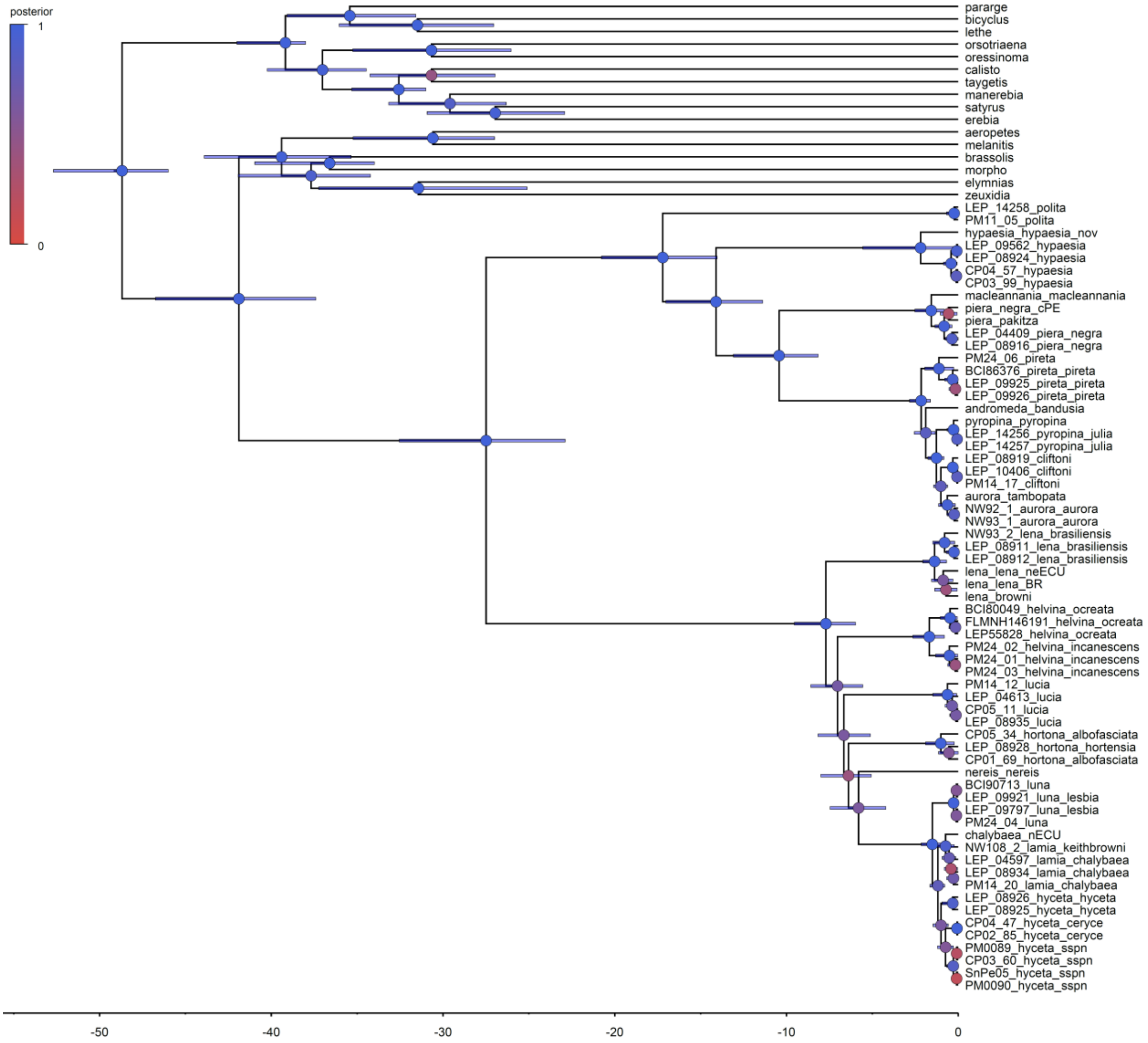

F: BP&P's delimitation model under $\theta$ = IG[3, 0.2] (large pop size)

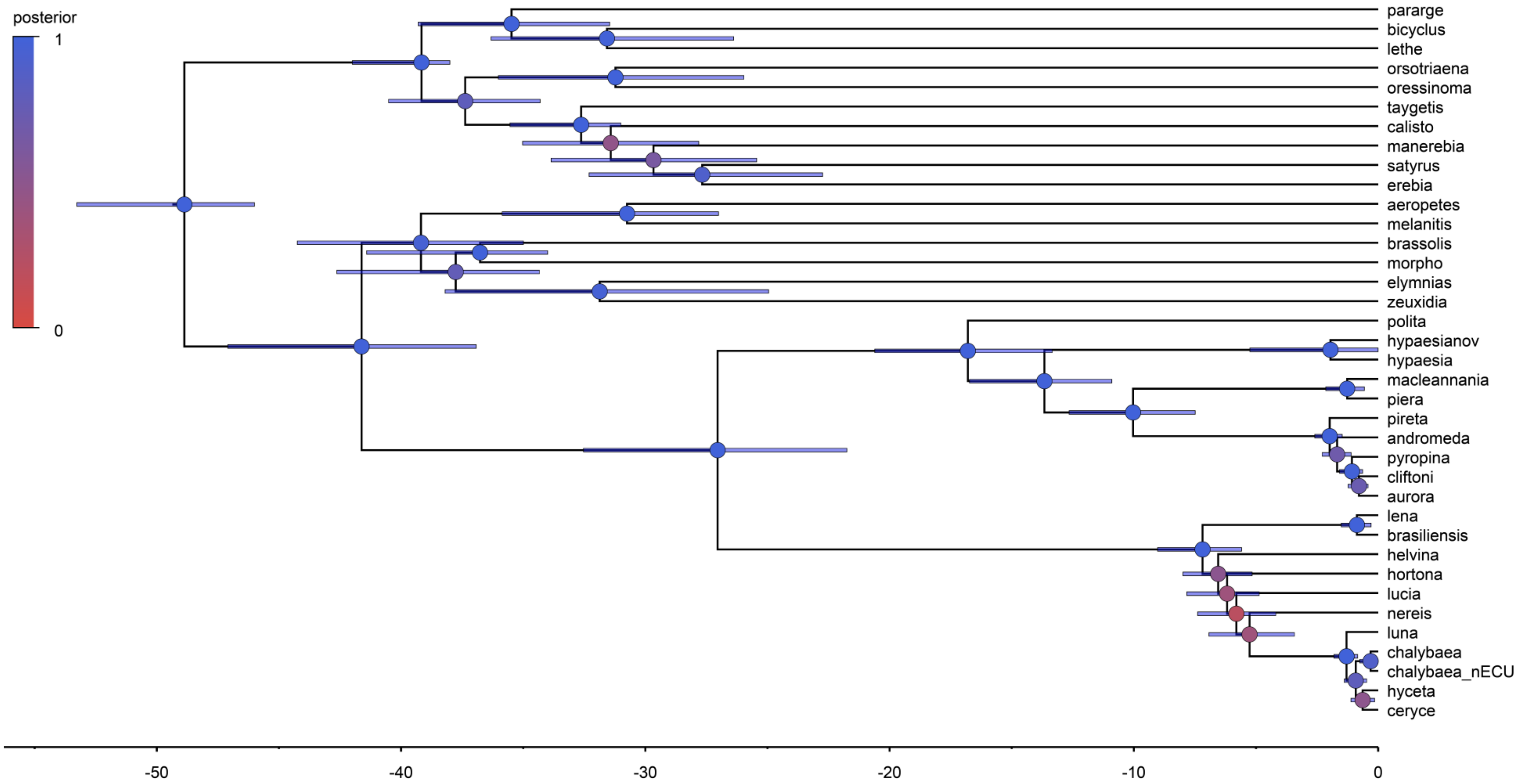

G: BP&P's delimitation model under θ = IG[3, 0.1] (medium pop size)

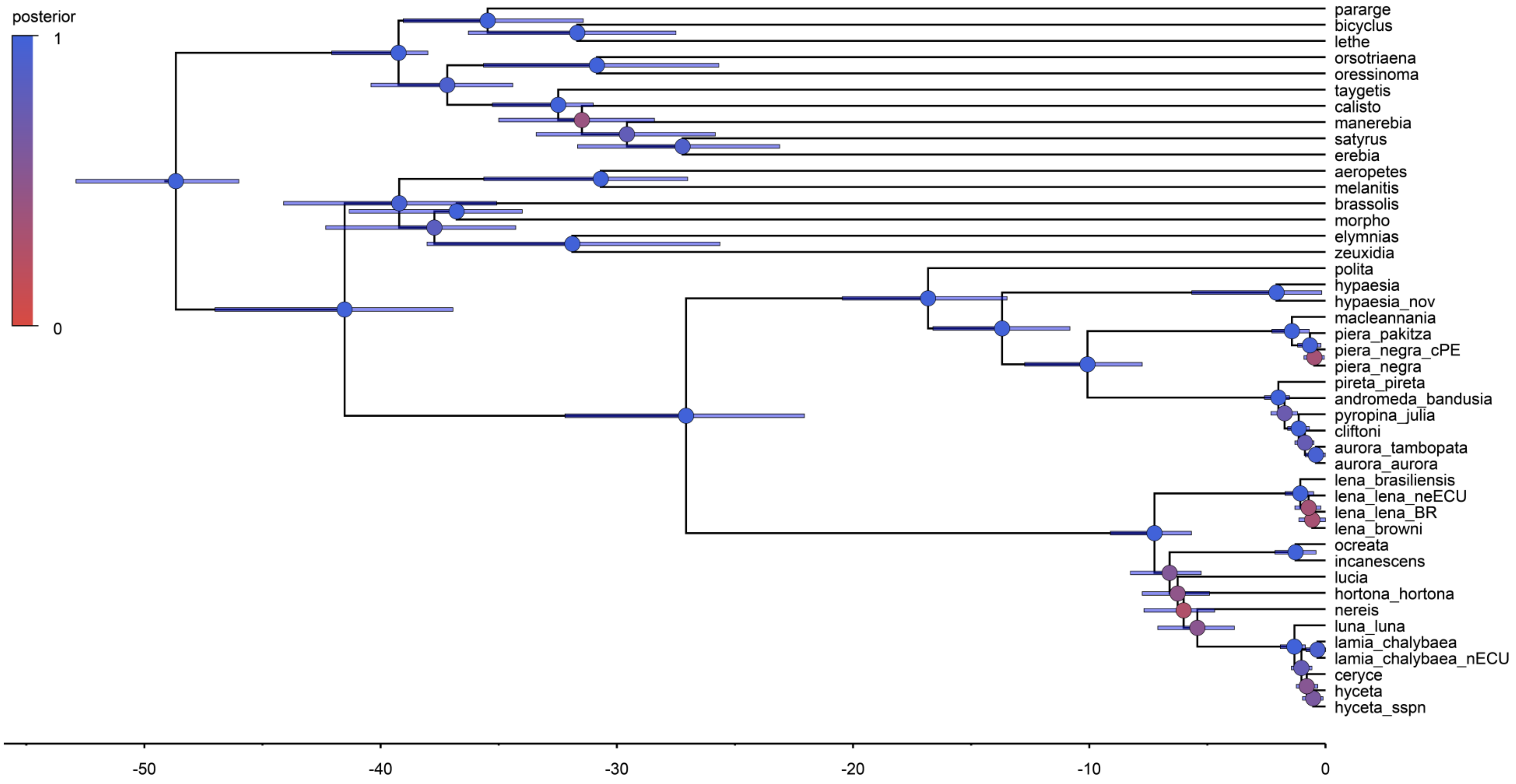

H: BP&P's delimitation model under $\theta$ = IG[3, 0.1] (small pop size)

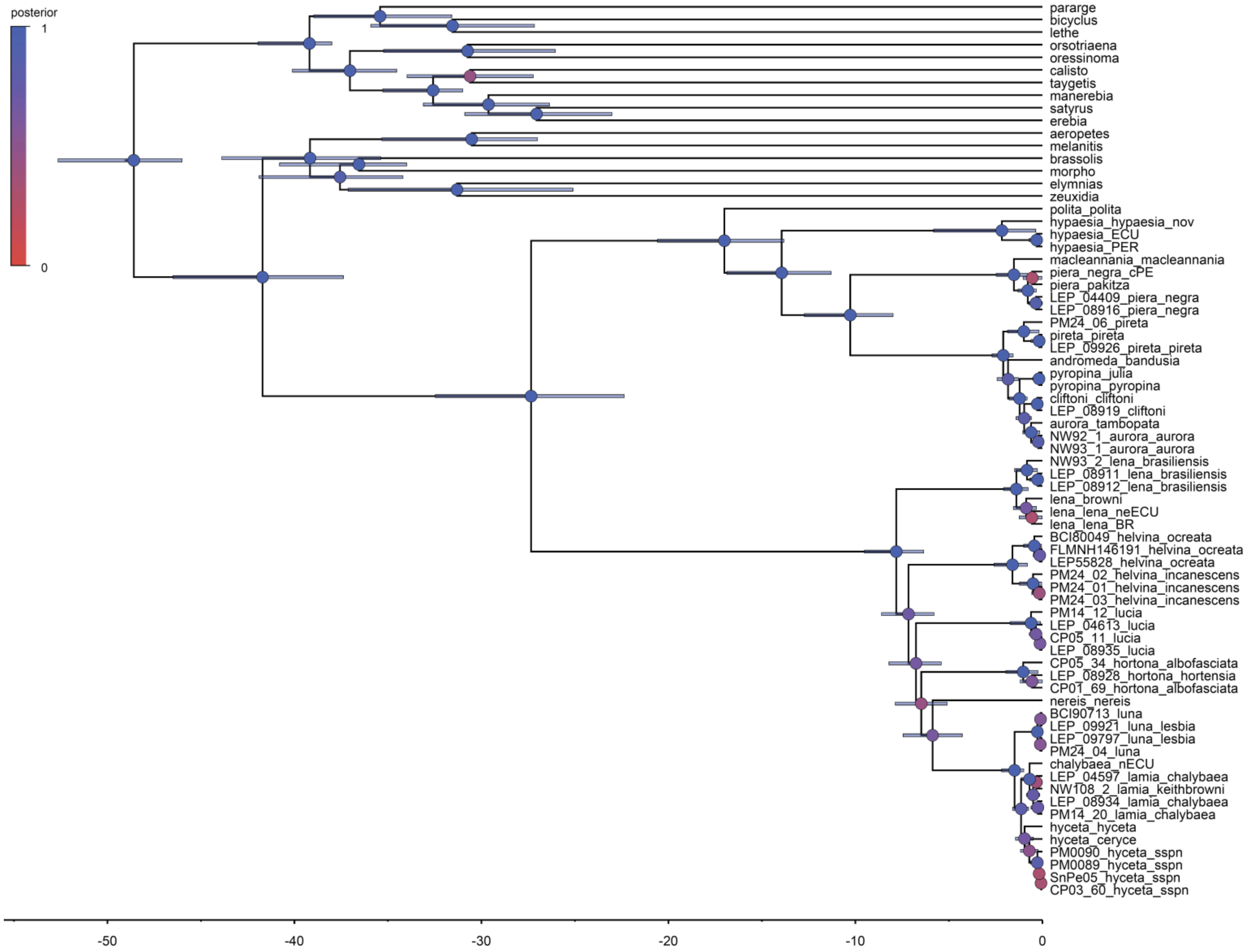
posterior
1
0
pararge
bicyclus
lethe
orsotriaena
oressinoma
calisto
taygetis
manerebia
satyrus
erebia
aeropetes
melanitis
brassolis
morpho
elymnias
zeuxidia
polita_polita
hypaesia_hypaesia_nov
hypaesia_ECU
hypaesia_PER
macleannania_macleannania
piera_negra_cPE
piera_pakitza
LEP_04409_piera_negra
LEP_08916_piera_negra
PM24_06_pireta
pireta_pireta
LEP_09926_pireta_pireta
andromeda_bandusia
pyropina_julia
pyropina_pyropina
cliftoni_cliftoni
LEP_08919_cliftoni
aurora_tambopata
NW92_1_aurora_aurora
NW93_1_aurora_aurora
NW93_2_lena_brasiliensis
LEP_08911_lena_brasiliensis
LEP_08912_lena_brasiliensis
lena_browni
lena_lena_neECU
lena_lena_BR
BCI80049_helvina_ocreata
FLMNH146191_helvina_ocreata
LEP55828_helvina_ocreata
PM24_02_helvina_incanescens
PM24_01_helvina_incanescens
PM24_03_helvina_incanescens
PM14_12_lucia
LEP_04613_lucia
CP05_11_lucia
LEP_08935_lucia
CP05_34_hortona_albofasciata
LEP_08928_hortona_hortensia
CP01_69_hortona_albofasciata
nereis_nereis
BCI90713_luna
LEP_09921_luna_lesbia
LEP_09797_luna_lesbia
PM24_04_luna
chalybaea_nECU
LEP_04597_lamia_chalybaea
NW108_2_lamia_keithbrowni
LEP_08934_lamia_chalybaea
PM14_20_lamia_chalybaea
hyceta_hyceta
hyceta_ceryce
PM0090_hyceta_sspn
PM0089_hyceta_sspn
SnPe05_hyceta_sspn
CP03_60_hyceta_sspn
-50
-40
-30
-20
-10
0
